# Parallel streams of raphe VGLUT3-positive inputs target the dorsal and ventral hippocampus in each hemisphere

**DOI:** 10.1101/2022.08.29.505760

**Authors:** Justine Fortin-Houde, Fiona Henderson, Guillaume Ducharme, Bénédicte Amilhon

**Author notes:** **Corresponding author:** Bénédicte Amilhon, Université de Montréal, Département de neurosciences, CHU Sainte-Justine Research Center, Montréal (QC) Canada.

## Abstract

The hippocampus (HP) receives neurochemically diverse inputs from the raphe nuclei, including glutamatergic fibers characterized by the expression of the vesicular glutamate transporter VGLUT3. These raphe-HP VGLUT3 (VGLUT3^HP^) projections have been suggested to play a critical role in HP functions, yet a complete anatomical overview of raphe VGLUT3 projections to the forebrain, and in particular the HP, is lacking. Using anterograde viral tracing, we describe largely non-overlapping VGLUT3-positive projections from the dorsal raphe (DR) and median raphe (MnR) to the forebrain, with the HP receiving inputs from the MnR. A limited subset of forebrain regions such as the amygdaloid complex, claustrum and hypothalamus receive projections from both the DR and MnR that remain largely segregated. This highly complementary anatomical pattern suggests contrasting roles for DR and MnR VGLUT3 neurons. To further analyse the topography of VGLUT3 raphe projections to the HP, we used retrograde tracing and found that VGLUT3^HP^ neurons distribute over several raphe sub-regions (including the MnR, paramedian raphe and B9 nucleus) and lack co-expression of serotonergic markers. Strikingly, two-color retrograde tracing unraveled two parallel streams of VGLUT3-positive projections targeting the dorsal and ventral poles of the HP. These results demonstrate highly organized and segregated VGLUT3-positive projections to the HP, suggesting independent modulation of HP functions such as spatial memory and emotion-related behavior.

## Introduction

Divisions along the longitudinal axis of the hippocampus (HP) are associated with distinct functions (Bannerman et al., 2004; Bannerman et al., 2014; Fanselow & Dong, 2010; Moser & Moser, 1998; Strange, Witter, Lein, & Moser, 2014). The dorsal part of the hippocampus (dHP) plays a crucial role in spatial navigation and memory (de Lavilleon, Lacroix, Rondi-Reig, & Benchenane, 2015; Moser, Moser, Forrest, Andersen, & Morris, 1995; Pothuizen, Zhang, Jongen-Relo, Feldon, & Yee, 2004) while the ventral part of the hippocampus (vHP) is essential to anxiety (Adhikari, Topiwala, & Gordon, 2010; Felix-Ortiz et al., 2013; Jimenez et al., 2018; Kjelstrup et al., 2002), social memory (Okuyama, Kitamura, Roy, Itohara, & Tonegawa, 2016; Rao, von Heimendahl, Bahr, & Brecht, 2019) and attention (Li et al., 2018; Yoshida et al., 2021). HP functional heterogeneity can be partly linked to topographically organized multimodal inputs from multiple brain regions, including inputs from the raphe nuclei (Muzerelle, Scotto-Lomassese, Bernard, Soiza-Reilly, & Gaspar, 2014; Vertes, Fortin, & Crane, 1999).

Hindbrain raphe nuclei (B1-B9) contain a mixed neuronal population including serotonergic (5-HT), GABAergic, dopaminergic and glutamatergic neurons (Fu et al., 2010; K. W. Huang et al., 2019; Sos et al., 2017; Szonyi et al., 2019). Long-range projecting glutamatergic neurons expressing the vesicular glutamate transporter type 3 (VGLUT3) are found in both the dorsal raphe (DR, corresponding to B6 and B7 nuclei) and the median raphe (MnR, corresponding to B5 and B8), and often co-express serotonergic neuronal markers (Fremeau et al., 2002; Gras et al., 2002; Herzog et al., 2004; Schafer, Varoqui, Defamie, Weihe, & Erickson, 2002). In the DR, VGLUT3-only and mixed VGLUT3/5-HT populations are mostly segregated. In the ventral part of the DR (DRV or B7v), VGLUT3-positive neurons colocalize extensively with 5-HT (Okaty et al., 2020; Ren et al., 2019) while in the dorsal part of the DR (DRD or B7d) a population of purely VGLUT3-positive neurons has been described (Hioki et al., 2010). DR VGLUT3 neurons send mixed 5-HT/glutamate projections to the amygdala where they can release glutamate in addition to 5-HT, although the contribution of glutamate to amygdala function remains unclear (Sengupta, Bocchio, Bannerman, Sharp, & Capogna, 2017; Sengupta & Holmes, 2019). In the ventral tegmental area and the orbitofrontal cortex, glutamate released by DR 5-HT neurons was suggested to contribute to vulnerability to social stress and active coping, respectively (Ren et al., 2018; Zou et al., 2020). Other studies focusing on DR VGLUT3 neurons have shown a role in reward and feeding behaviors (Liu et al., 2014; McDevitt et al., 2014; Nectow et al., 2017; Qi et al., 2014; H. L. Wang et al., 2019).

In the MnR, VGLUT3-positive neurons represent approximately 26% of the total MnR neuron population; overlap with 5-HT markers is limited since only 13% of all MnR VGLUT3 neurons co-express 5-HT (Sos et al., 2017). MnR VGLUT3 neurons project to the prefrontal cortex, medial septum and dHP (Jackson, Bland, & Antle, 2009; Szonyi et al., 2016; Varga et al., 2009) and occasionally send collaterals to several regions (Szonyi et al., 2016). So far, MnR VGLUT3-positive projections have been mostly studied in the HP, where they mediate fast excitatory transmission onto interneurons (He et al., 2022; Varga et al., 2009) and have been suggested to underlie the powerful control of the MnR on dHP activity, especially theta rhythm (Crooks, Jackson, & Bland, 2012; Domonkos et al., 2016; W. Huang, Ikemoto, & Wang, 2022).

Despite tantalizing evidence for their role in modulating HP function and rhythms, the topographical organization and properties of raphe-HP VGLUT3 projections are unknown. More broadly, forebrain glutamatergic projections from the DR and MnR have been partially described but no comprehensive comparative overview is available to date (Geisler, Derst, Veh, & Zahm, 2007; He et al., 2022; Jackson et al., 2009; Liu et al., 2014; McDevitt et al., 2014; Nectow et al., 2017; Qi et al., 2014; Ren et al., 2018; Senft, Freret, Sturrock, & Dymecki, 2021; Sengupta et al., 2017; Sengupta & Holmes, 2019; Szonyi et al., 2016; Varga et al., 2009; H. L. Wang et al., 2019; Zou et al., 2020). In this study, we systematically compare DR versus MnR VGLUT3-positive projections to the forebrain, with a focus on their projections to the HP. We describe non-overlapping DR versus MnR VGLUT3-positive projections to the forebrain, with the HP showing innervation predominantly by the MnR. To precisely pinpoint the origins of raphe-HP glutamatergic input we turned to retrograde labelling to map HP-projecting VGLUT3-positive (VGLUT3^HP^) neurons within raphe nuclei. We find that VGLUT3^HP^ neurons are located in several subregions of the ventral part of the raphe and show little co-expression with 5-HT. We uncover distinct glutamatergic populations unilaterally targeting the dHP and the vHP and describe their topographical organization in both the medio-lateral and rostro-caudal axis. Our results extend previous knowledge regarding VGLUT3-positive projections from the raphe and provide the first in-depth anatomical characterization of the raphe-HP glutamatergic pathway. Notably, we reveal parallel streams of glutamatergic projections to the dHP and vHP, suggesting differential modulation of HP function over the longitudinal axis.

## Materials and Methods

### Animals

All experiments were conducted in accordance with the Canadian Council on Animal Care Guidelines and were approved by the Comité Institutionnel de Bonnes Pratiques Animales en Recherche (CIBPAR) at Sainte-Justine Hospital research center. Mice were group-housed in a temperature- and humidity-controlled room on a 12/12-hr light/dark cycle with *ad libitum* access to food and water. Adult (2-4 months old) male (n = 12) and female (n = 8) VGLUT3-Cre mice were used for experiments; the VGLUT3-Cre mouse strain was a generous gift by Dr El Mestikawy (Centre de Recherche de l’Hôpital Douglas, Québec, Canada). All VGLUT3-Cre mice were heterozygotes and maintained on a C57BL6/J background. In addition, wild-type C57BL6/J adult male (n = 2) and female (n = 2) mice were used to control for non-specific expression of Cre-dependent adeno-associated viruses (AAVs).

### Double-probe fluorescent in situ hybridisation (sdFISH)

sdFISH was carried out as described previously (Dumas & Wallen-Mackenzie, 2019). sdFISH was performed using antisense riboprobes for the detection of Slc17a8 mRNA: NM_182959.3 sequence 1902-2320 and Cre: AB449974.1 sequence 245-1060. Synthesis of digoxigenin and fluorescein-labeled RNA probes were made by a transcriptional reaction with incorporation of digoxigenin or fluorescein labelled nucleotides. Specificity of probes was verified using NCBI blast. Cryosections were air-dried, fixed in 4% paraformaldehyde (PFA) and acetylated in 0.25% acetic anhydride/100 mM triethanolamine (pH 8). Sections were hybridized for 18 h at 65 °C in 100 µl of formamide-buffer containing 1 µg/ml digoxigenin-labeled riboprobe (DIG) and 1 µg/ml fluorescein-labeled riboprobe. Sections were washed at 65 °C with SSC buffers of decreasing strength and blocked with 20% fetal bovine serum and 1% blocking solution. Fluorescein epitopes were detected with horseradish peroxidase (HRP) conjugated anti-fluorescein antibody at 1:5000 and revealed using Cy2-tyramide at 1:250. HRP-activity was stopped by incubation of sections in 0.1 M glycine followed by a 3% H_2_O_2_ treatment. DIG epitopes were detected with HRP anti-DIG Fab fragments at 1:2000 and revealed using Cy3 tyramide at 1:100. Nuclear staining was performed with 4’ 6-diamidino-2-phenylindole (DAPI).

### Viral vectors

AAVdj-EF1a-DIO-hChR2_(E123T/T159C)_-eYFP viral vector (6.5E12 GC/ml) produced by the Canadian Neurophotonics Platform Viral Vector Core Facility (RRID:SCR_016477) was used for anterograde tracing experiments. Retrograde tracing was performed using the viral vectors AAV2retro-CAG-FLEX-tdTomato (1.3E13 GC/ml) obtained from Addgene (provided by Edward Boyden; Addgene plasmid#28306; http://n2t.net/addgene:28306; RRID:Addgene_28306) and AAV2retro-Ef1a-DIO-eYFP (5.5E12 GC/ml) obtained from the Canadian Neurophotonics Platform Viral Vector Core Facility (RRID:SCR_016477).

### Stereotaxic surgeries/injections

Anesthesia was induced using 5% isoflurane and maintained with 1% isoflurane during surgery. Body temperature was maintained at 37 °C through an electric heating pad and carprofen (2 mg/kg, subcutaneous) was administered before the surgery. Mice were secured in the stereotaxic apparatus and craniotomies were performed over the raphe nuclei (for anterograde tracing) or the HP (for retrograde tracing). Glass micropipettes were back filled with mineral oil and loaded with viral vectors. Injections were done at 1 nL/s with a Nanoject III (Drummond Scientific Company).

For DR anterograde tracing (Figure 1 and 4), a total volume of 200 nL (n = 1) or 500 nL (n = 2) was injected over two sites at the following coordinates (in mm relative to bregma): anterior-posterior (AP) -4.36 and -4.80, medio-lateral (ML) 0, dorso-ventral (DV) -3.00. For MnR anterograde tracing (Figures 2-4), a volume of 25 nL (n = 1) or 35 nL (n = 1) was injected at coordinates: AP -4.23 or -4.48, ML 0 and DV -4.80. For all raphe injections, the glass capillary was inserted with a 10° lateral angle to avoid damaging the confluence of sagittal and transverse sinus. For retrograde tracing of VGLUT3-positive raphe inputs to the entire HP (Figure 5), 800 nL of AAV2retro-CAG-FLEX-tdTomato was injected bilaterally in the dHP at AP -2.30, ML ±1.35, DV -1.62 and 400 nL of AAV2retro-Ef1a-DIO-eYFP was injected bilaterally in the vHP AP -3.65, ML ±3.25, DV -4.00. For retrograde tracing of inputs to the dHP versus vHP (Figure 6), the same coordinates were used with smaller injection volumes: 400 nL (n = 4) of AAV2retro-CAG-FLEX-tdTomato was injected bilaterally in the dHP (n = 3) or vHP (n = 1) and 300 nL of AAV2retro-Ef1a-DIO-eYFP was injected bilaterally in the dHP (n = 1) or vHP (n = 3). In a second set of experiments (“small volumes” Figure 5E), viral vector volumes were further decreased to fully avoid overlap between dHP- and vHP-targeting injections. Two mice received a volume of 200 nL of AAV2retro-Ef1a-DIO-eYFP bilaterally in the dHP and 200 nL of AAV2retro-CAG-FLEX-tdTomato bilaterally in the vHP. Finally, for retrograde tracing of VGLUT3 raphe inputs to the right versus left HP (Figure 7), dHP coordinates were moved to AP -2.30, ML ±2, DV -1.62, an intermediate injection site was added at AP: -2.92, ML: ±2.7, DV: -2.3 and the vHP site remained the same. A volume of 300 nL of AAV2retro-Ef1a-DIO-eYFP or AAV2retro-CAG-FLEX-tdTomato was injected at each site in right or left HP respectively. To control for non-specific expression of the viral vectors, two C57BL6/J mice were injected in the dHP (300 nL) and vHP (400 nL) with AAV2retro-Ef1a-DIO-eYFP and two C57BL6/J mice injected in the dHP (400 nL) and vHP (600 nL) with AAV2retro-CAG-FLEX-tdTomato.

**Figure 1.**
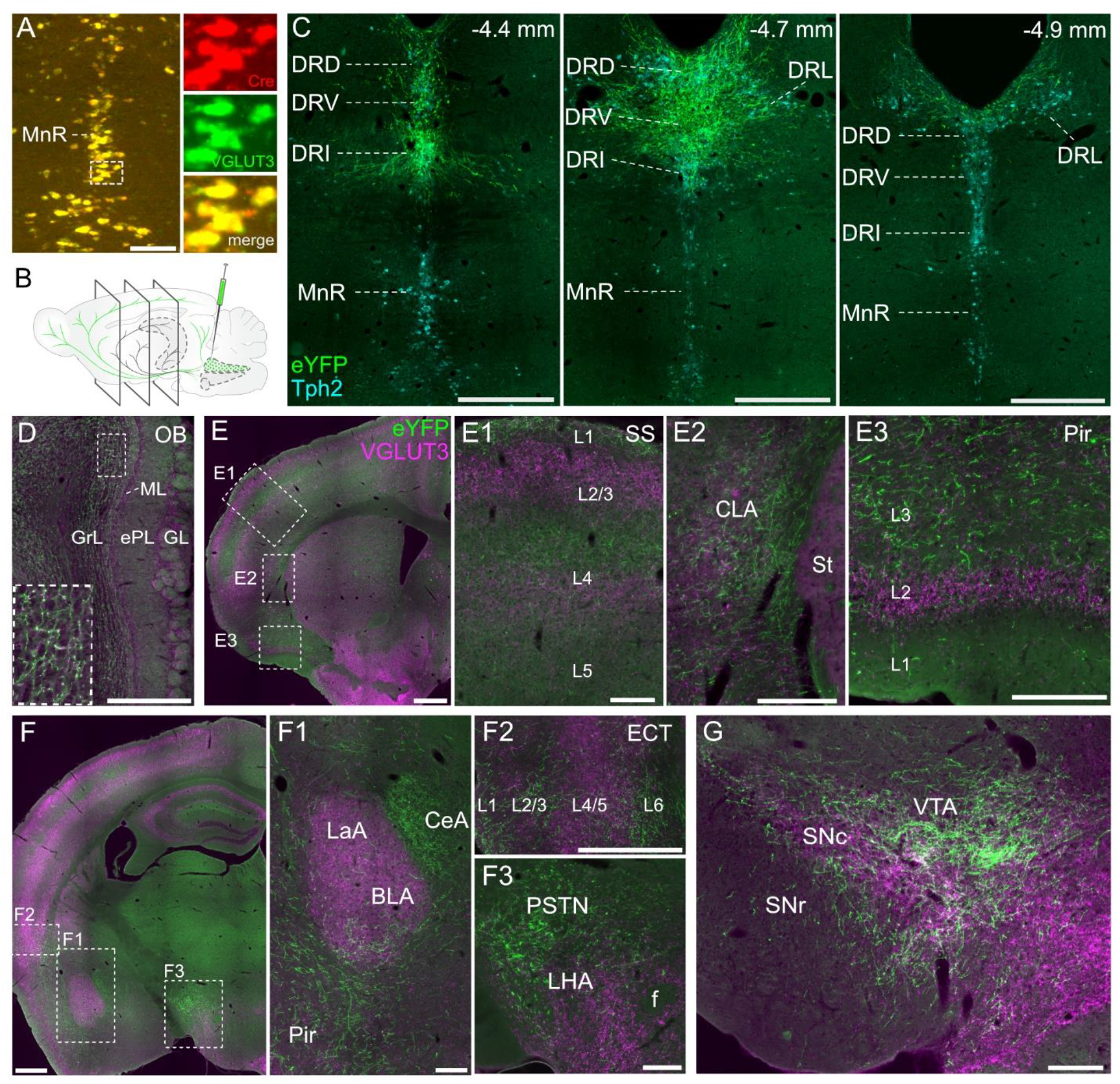
Dorsal raphe VGLUT3 projections to the forebrain. (**A**) Validation of Cre expression specificity in VGLUT3-Cre mice using double fluorescent *in situ* hybridization detecting mRNAs for VGLUT3 (green) and Cre (red). (**B**) Schematic of Cre-dependent viral vector injection in the DR of VGLUT3-Cre mice. (**C**) Representative eYFP expression (green) in the DR at different rostro-caudal levels (distance is indicated in mm from bregma) with large coverage over the rostro-caudal axis, showing expression in all sub-regions of dorsal raphe (DR), including dorsal (DRD), ventral (DRV), interfascicular (DRI) and lateral (DRL). Identification of raphe subdivisions using Tph2 immunofluoresence (cyan) shows no leakage in MnR. (D-G) Coronal sections of forebrain regions showing dense eYFP-positive fibers expression detected using anti-GFP immunostaining. All sections are co-stained for VGLUT3 (magenta). (**D**) In the olfactory bulb (OB) a high density of fibers is observed in the granular cell layer (GrL, *inset*), with very few fibers in the glomerular layer (GL), external plexiform layer (ePL) and mitral cell layer (ML). (**E**) High fiber density is found in the somatosensory cortex (SS, E1) with layers 1 to 5 indicated (L1-L5), the claustrum (CLA, E2) closely apposed to the striatum (St) and the piriform cortex (Pir, E3) mainly in layer 3. (**F**) At more rostral levels, variable density of DR VGLUT3 innervation can be observed in the amygdala, with a high density of fibers in the central amygdala (CeA), lower density in the basolateral amygdala (BLA) and sparse fibers in the lateral amygdala (LaA). Dense DR glutamatergic inputs are also present in the ectorhinal cortex (ECT, F2), parasubthalamic nucleus (PSTN) and lateral hypothalamic area (LHA, F3), lateral to the fornix (f). (**G**) A high density of fibers was also observed in the ventral tegmental area (VTA) with some fibers present in the substantia nigra pars compacta (SNc) and the medio-dorsal substantia nigra pars reticulata (SNr). Scale bars: 500 µm in C, D, E, F and G, 150 µm in A, E1-3 and F1-3.

**Figure 2.**
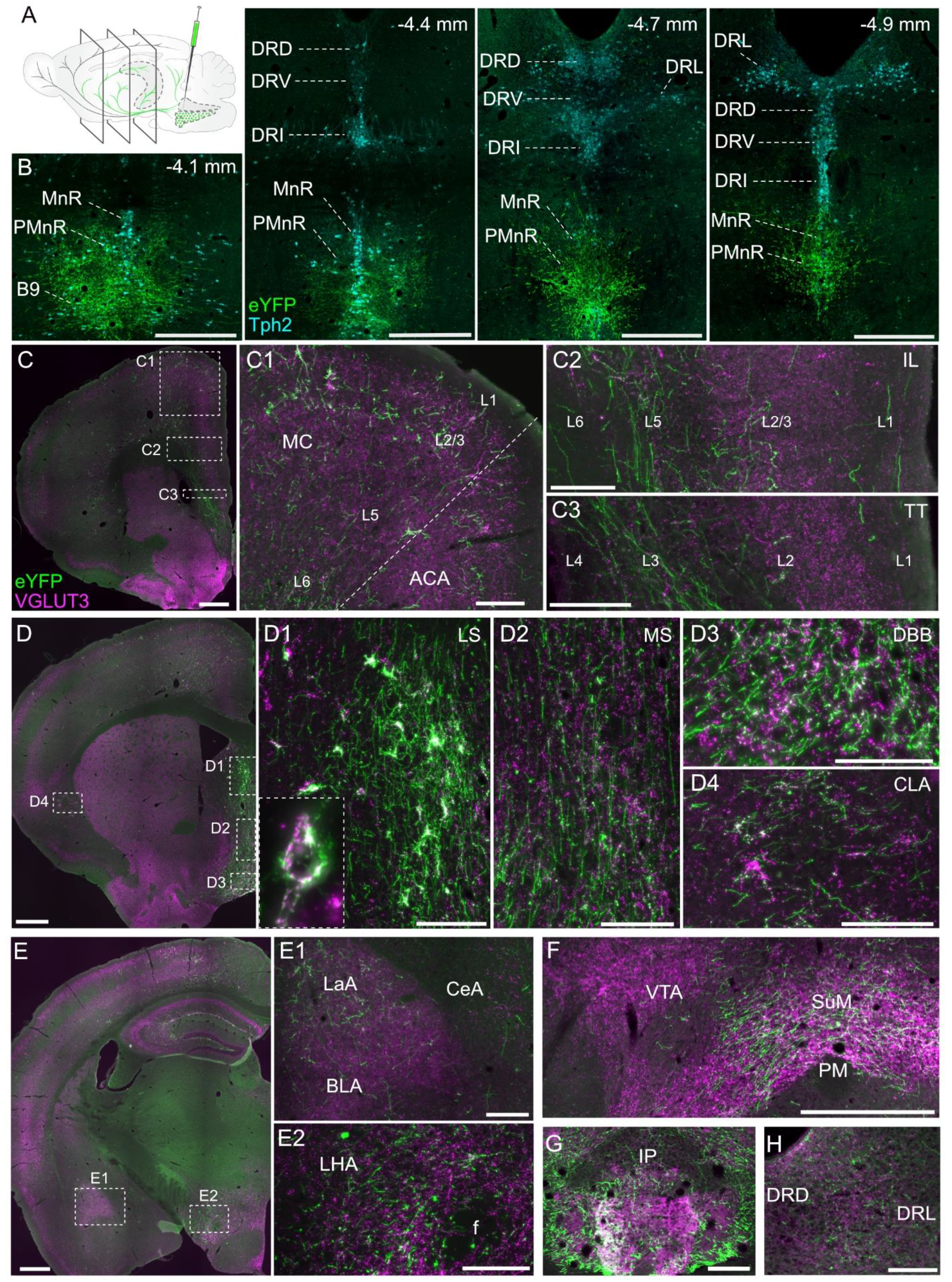
Median raphe region VGLUT3 projections to the forebrain. (**A**) Schematic of Credependent viral vector injection in the MnR region of VGLUT3-Cre mice. (**B**) Representative eYFP expression (green) in the MnR region at different raphe levels (distance in mm from bregma) showing large coverage over the rostro-caudal axis in the MnR, paramedian (PMnR) parts of the region and B9 nucleus and absence of expression in the DR. (C-G) Coronal sections of the forebrain regions showing anti-GFP (green) and VGLUT3 (magenta) immunostainings. (**C**) In frontal regions, highest density of eYFP fibers is detected in the motor cortex (MC, C1) especially in L2/3 and L6, anterior cingulate area (ACA), layer 5 of prelimbic (PL) and infralimbic (IL) cortices (C2) and layer 3 of teania tecta (TT, C3). (**D**) Dense MnR region inputs are found in the lateral septum (LS, D1, *inset* on eYFP terminals forming a pericellular basket was taken from another section), medial septum (MS, D2) and diagonal band of Broca (DBB, D3). Glutamatergic fibers can also be observed in the ventral part of the claustrum (CLA, D4). (**E**) In the amygdala, a high density of fibers can be seen in the lateral amygdala (LaA), sparse expression in the basolateral amygdala (BLA) and no expression in the central amygdala (CeA, E1). Dense fibers are present in the lateral hypothalamic area (LHA, E2) adjacent to the fornix (f). (**F**) Supramammillary nucleus (SuM) located over the principal mammillary tract (PM) was heavily innervated while VTA contained almost no fibers. (**G**) The interpeduncular nucleus (IP) showed high density of glutamatergic fibers. (**H**) The DR receives VGLUT3 input from the MnR in the DRD and DRL regions. Scale bars: 500 µm in B, C, D, E and F, 150 µm in C1-3, D1-4, E1-2, G and H.

**Figure 3.**
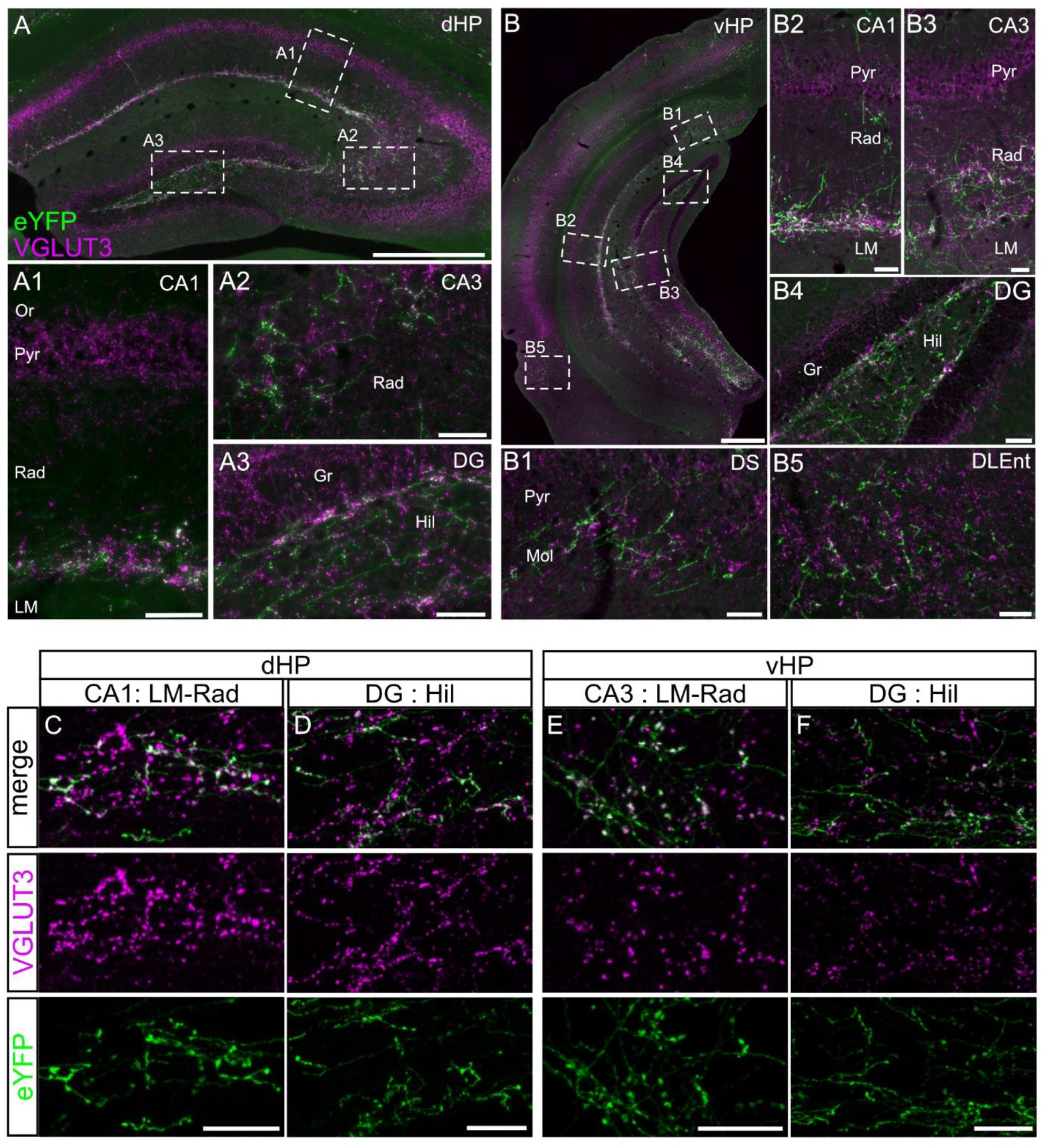
Median raphe region VGLUT3 projections to the hippocampus. (**A**) In the dorsal hippocampus (dHP), VGLUT3 inputs from the MRR were found in CA1 (A1) at the interface between stratum radiatum (Rad) and stratum lacunosum-moleculare (LM). Sparse fibers were observed in the stratum oriens (Or) and virtually no fibers were found in the pyramidal layer (Pyr). In the CA3 (A2), MRR VGLUT3 fibers were dense in stratum radiatum. In the dentate gyrus (DG, A3), glutamatergic fibers were present in the hilus (Hil) but sparse or absent in the granular layer (Gr). (**B**) A similar pattern of expression was observed in the ventral HP (vHP) across CA1 (B2), CA3 (B3) and DG (B4). In addition, we noted moderate to high fiber density in the molecular layer of the subiculum (SUB, B1) and dorsolateral entorhinal cortex (DLEnt, B5). (**C-F**) A high degree of colocalization between eYFP and VGLUT3 was observed across all subregions of the HP, and other brain areas (not shown). Scale bars: 500 µm in A and B, 150 µm in A1-3 and B1-5, 20 µm in C-F.

**Figure 4.**
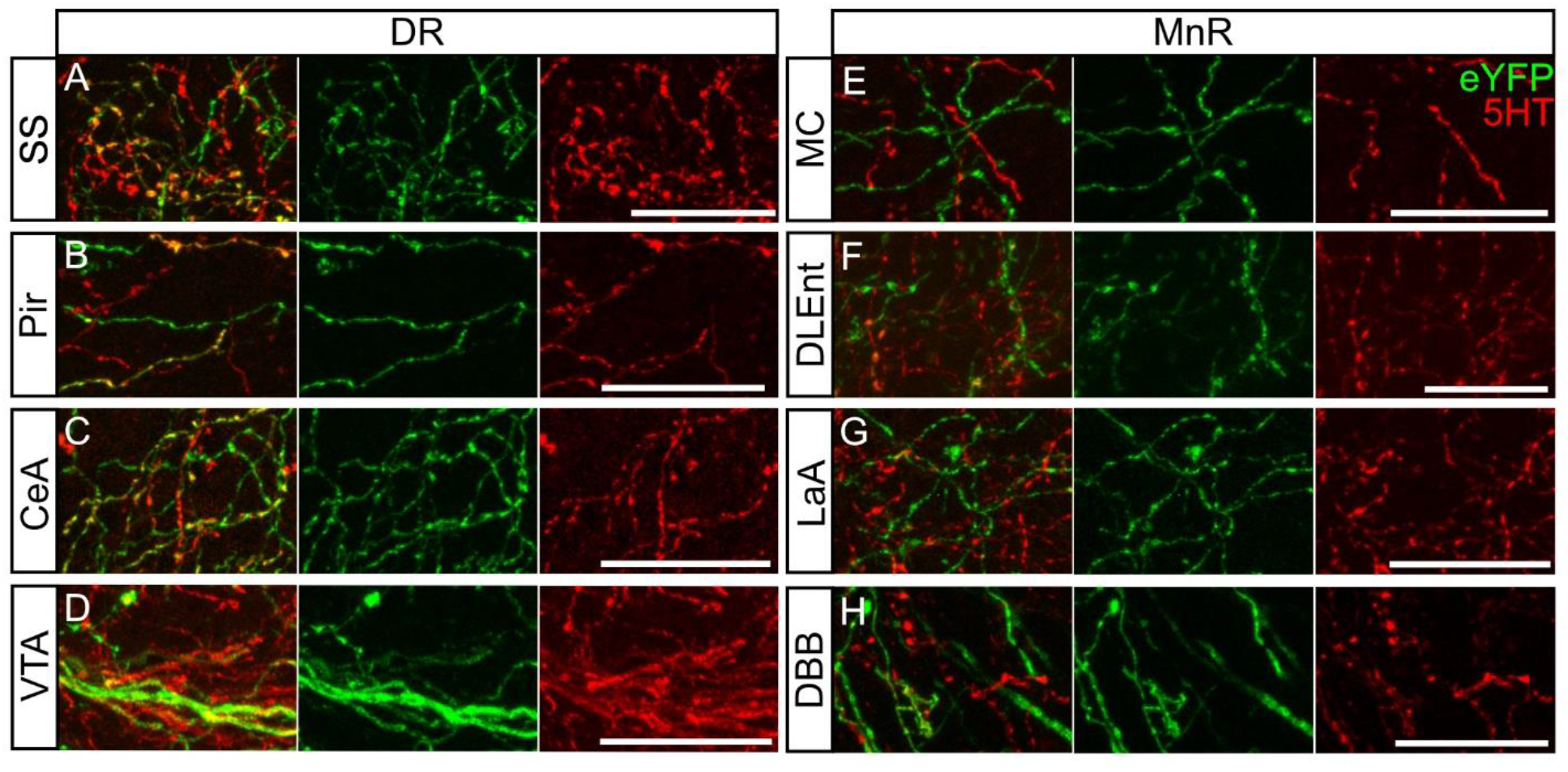
Co-expression of serotonin (5-HT) in eYFP fibers from the DR and MnR. Colocalization between eYFP in VGLUT3-positive fibers (green) and 5-HT (red) was assessed qualitatively. Colocalization between eYFP and 5-HT was frequently observed in VGLUT3-positive fibers arising from the DR (left), as exemplified in the SS cortex (**A**), Pir cortex (**B**), CeA (**C**) and VTA (**D**). In contrast, VGLUT3-positive fibers coming from the MnR (right) showed little or no colocalization with 5-HT, see for example MC (**E**), DLEnt cortex (F), LaA (G) and DBB (H). Scale bar: 20 μm.

**Figure 5.**
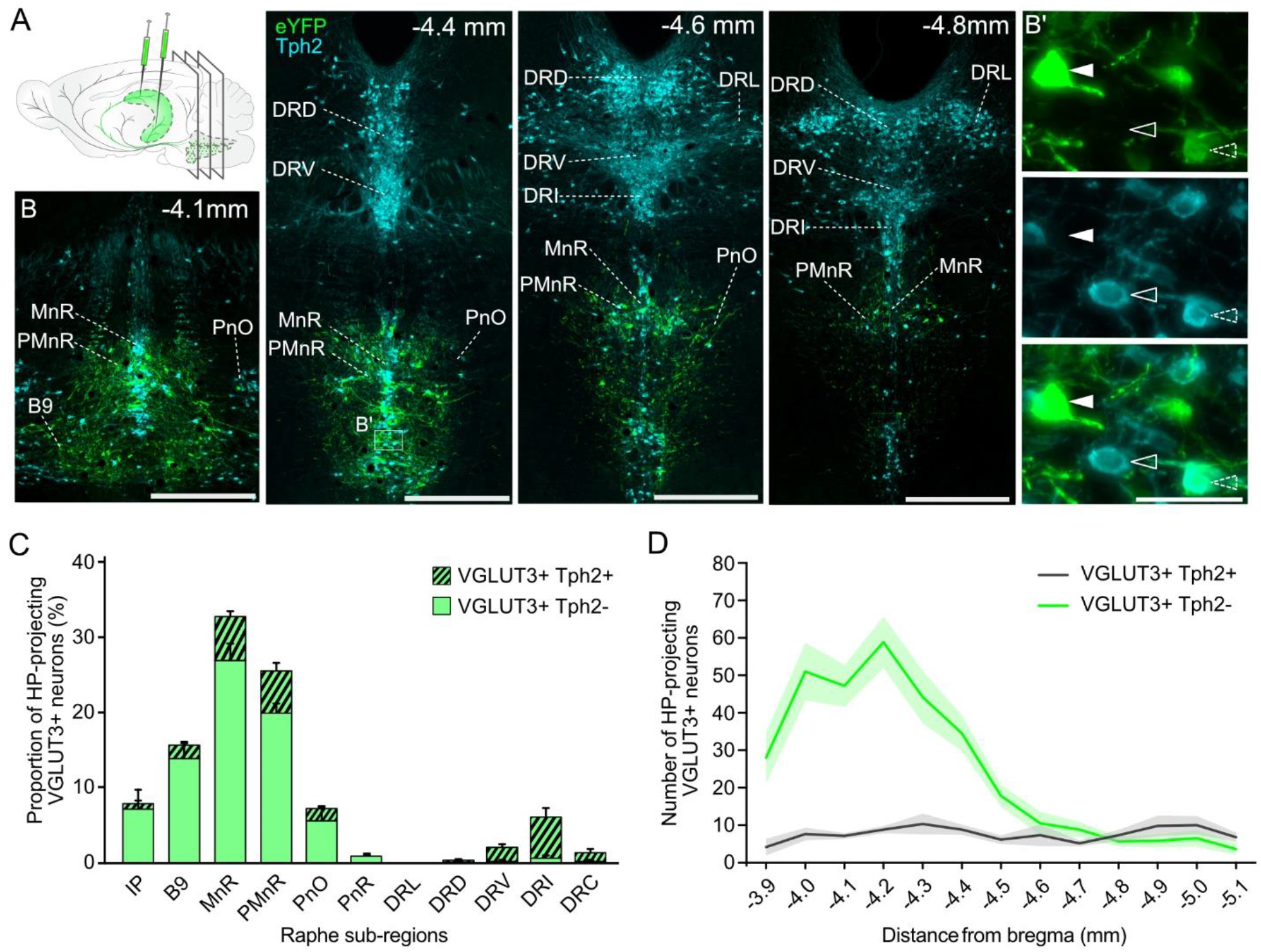
Hippocampus-projecting VGLUT3-positive (VGLUT3^HP^) neurons distribute over multiple raphe nuclei. (**A**) Schematic of Cre-dependent retrograde viral vector injection in dorsal and ventral parts of the hippocampus of VGLUT3-Cre mice. Injections were bilateral for all mice. (**B**) Representative eYFP expression (green) in the raphe at different levels (in mm from bregma). Overall, VGLUT3^HP^ somas were largely restricted to the ventral part of the raphe nuclei. Co-expression of serotonin was determined based on anti-Tph2 immunostaining (cyan). Inset in (B’) shows a Tph2-negative VGLUT3^HP^ neuron (full arrow), a Tph2-postive VGLUT3^HP^ neuron (dashed arrow) and an eYFP-negative 5-HT neuron (empty arrow). (**C**) Distribution of VGLUT3+Tph2-(green) and VGLUT3+Tph2+ (stripes) neurons in raphe subregion. Results are shown as the proportion of all VGLUT3^HP^ neurons (293-536 VGLUT3^HP^ neurons per mouse, n = 6 mice). VGLUT3^HP^ neurons are mainly located in the IP, B9 nucleus, PnO, MnR and PMnR. Note the low proportion of VGLUT3^HP^ neurons that co-express the serotonergic marker Tph2. PnR: pontine raphe nucleus. (**D**) Rostro-caudal distribution of VGLUT3+Tph2-neurons (green) and VGLUT3+Tph2+ neurons (grey). VGLUT3+Tph2-neurons are mainly located in the rostral half of the raphe while VGLUT3+Tph2+ are evenly distributed (n = 6 mice). Scale bars: 500 µm in B, 50 µm in B’.

**Figure 6.**
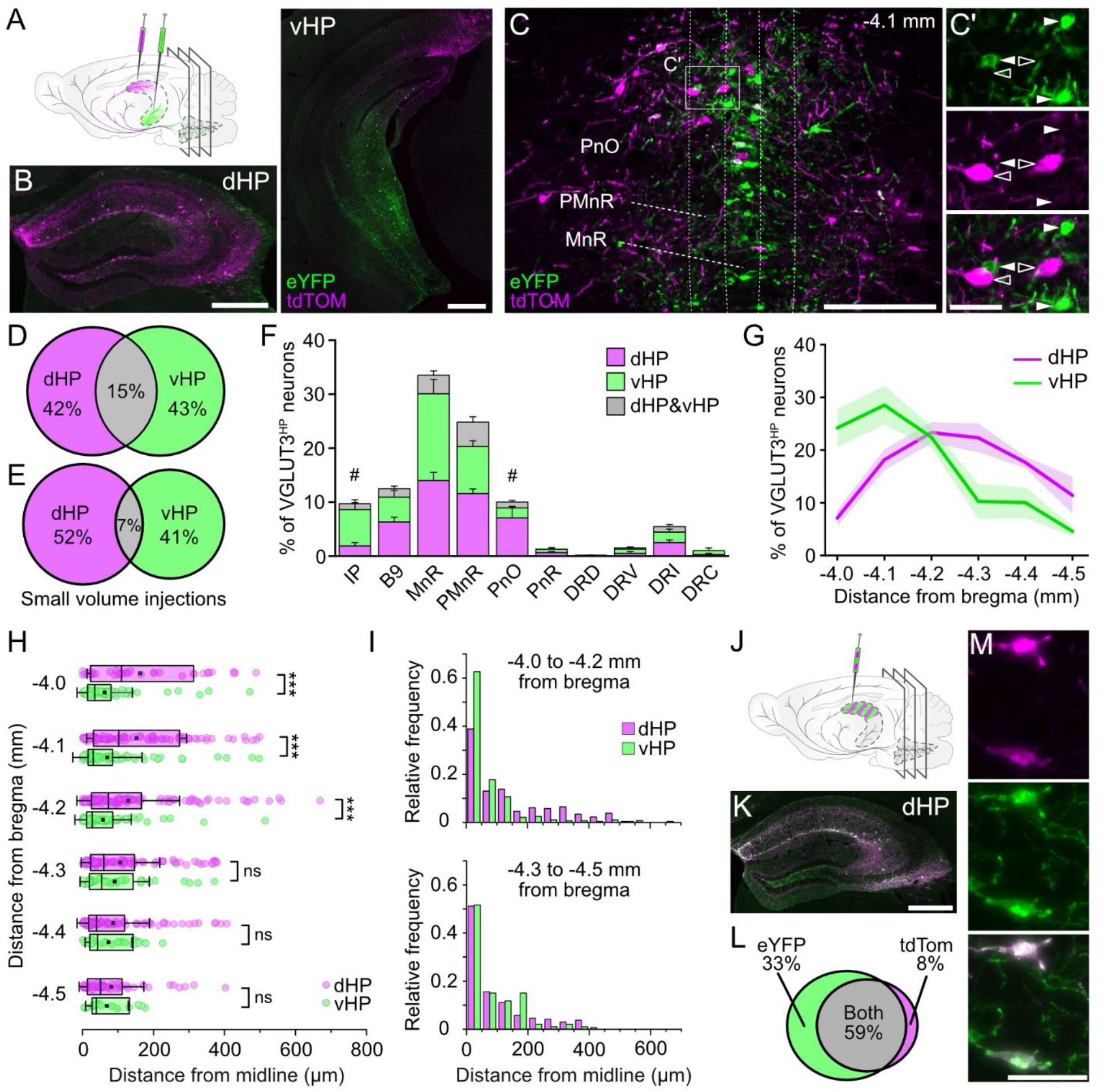
Separate populations of VGLUT3^HP^ neurons project to the dorsal and ventral parts of the hippocampus. (**A**) Schematic of two-color Cre-dependent retrograde viral vector injection in VGLUT3-Cre mice. Viral vectors expressing tdTomato (tdTOM) or eYFP were injected bilaterally in the dorsal hippocampus (dHP) and ventral hippocampus (vHP). (**B**) Representative tdTOM (magenta) and eYFP (green) expression dHP and vHP. (**C**) Representative tdTOM (magenta) and eYFP (green) retrograde viral expressions in the MRR at -4.1 mm from bregma. Inset (C’) shows dHP-projecting VGLUT3+ neurons (empty arrows) and vHP-projecting VGLUT3+ neurons (full arrows). (**D**) Venn diagram representing the proportions of dHP-, vHP- and double-projecting VGLUT3+ neurons (1648 neurons, n = 4 mice). (**E**) Venn diagram showing dHP-, vHP- and double-projecting VGLUT3+ neurons with small volumes injections to eliminate all overlap at the injection site (599 neurons, n = 2 mice). (**F**) Proportion of VGLUT3+ neurons projecting to the dHP, vHP or both per raphe region. Results are shown as the proportion of all VGLUT3^HP^ neurons (276-515 VGLUT3+ neurons per mouse, n = 6 mice). Equivalent proportions of dHP- and vHP-projecting VGLUT3+ neurons are found in all raphe subnuclei except in the IP and PnO. (**G**) Rostro-caudal distribution of dHP- and vHP-projecting VGLUT3+ neurons in the MRR (i.e. MnR, PMnR and PnO). vHP-projecting VGLUT3+ neurons are mainly located in the first half of the MRR while dHP-projecting VGLUT3+ neurons are more evenly distributed (n = 6 mice). (**H**) Spatial distribution of dHP- and vHP-projecting VGLUT3 neurons at different rostro-caudal levels of the MRR. Distance from midline was calculated for each individual dHP- and vHP-projecting VGLUT3 soma (circles). Box represents 25-75 percentile; line in the box is the median and square symbol is the mean; error bars represent 1 SD. In the rostral half of the MRR, dHP-projecting neurons are located significantly further from the midline. (**I**) Relative frequency of neurons in rostral (top) and caudal (bottom) parts of the MRR, bins size is 50 µm. (**J**) Schematic of the control experiment where a mix of the two Cre-dependent retrograde viral vectors was injected in the dHP. (**K**) Representative tdTOM (magenta) and eYFP (green) expression in the dHP. (**L**) Venn diagram showing VGLUT3+ neurons expressing eYFP, tdTOM or both among all dHP-projecting VGLUT3+ neurons in B9 and MRR (263 neurons, n = 2 mice). (**M**) Representative tdTOM (magenta) and eYFP (green) retrograde viral co-expression in two neurons of the MRR. Scale bar: 500 µm in B and K, 250 µm in C and 50 µm in C’ and M. ns: non significant; *** *p* < 0.001

### Tissue processing and immunofluorescence

Mice were deeply anesthetized with a mixture of ketamine (80 mg/kg), xylazine (12.5 mg/kg) and acepromazine (2.5 mg/kg) and transcardially perfused with 4% PFA in phosphate buffer saline (PBS). Brains were removed, incubated in 4% PFA for 24 h and cryoprotected using 15% sucrose in PBS (24 h). Brains were embedded in Tissue-Plus™ O.C.T. Compound (Fisher scientific cat. No 23-730-571) frozen in isopentane and stored at −80°C until sectioning. Free floating coronal sections (25 μm) were cut on a cryostat (Leica Biosystems model CM3050S) and stored in PBS-azide 0.02%. Sections were collected along the entire rostro-caudal axis of the brain. Sections were first incubated 3 × 45 min in a PBS-gelatin 0.45%-triton 0.25% blocking buffer (PGT). Sections were then incubated with the following primary antibodies for 48 h at 4 °C: chicken anti-GFP (1:1000, Invitrogen cat. No. A10262), rabbit anti-RFP (1:10000, VWR Rockland cat. No. CA600-401-379), guinea pig anti-Tph2 (1:1000, Frontier Institute cat. No. TPH2-GP-Af900) or guinea pig anti-VGLUT3 (1:1000, Synaptic Systems cat. No. 135204). Sections were then washed in PGT (3 × 15 min) and incubated in the following secondary antibodies for 1.5 h at room temperature: donkey anti chicken-A488 (1:2000, Jackson Immunoresearch cat. No. 703-545-155), donkey anti rabbit-A555 (1:2000, Invitrogen cat. No. A31572) and goat anti guinea pig-A647 (1:2000, Invitrogen cat. No. A21450). Sections were finally washed in PBS (3 × 15 min) and mounted on glass slides (Superfrost plus, Fisherbrand cat. No. 12-550-15) using a DAPI mounting medium (Fluoromount-G with DAPI, Invitrogen cat. No.00-4959-52).

### Image acquisition and data analysis

For sdFISH image acquisition, slides were scanned with a 20X objective using the NanoZoomer 2.0-HT (Hamamatsu, Japan). For tracing experiments, images were acquired using a Leica DMi8 wide-field inverted microscope with 20X and 40X objectives. Images were acquired in z-stacks (step size 2 µm) and cell counting was performed manually using the FIJI multi-point tool. The maximum projection feature of LasX software (Leica) was used to illustrate the distribution of fibers and somas in the various brain regions analyzed. Colocalization of the viral tracer with 5-HT or VGLUT3 immunostainings was assessed using a Leica SP8 confocal microscope with 63X objective (Figures 3 and 4). Pictures shown are maximum projections of one-µm step z-stacks.

Injection sites were systematically confirmed and only mice with viral expression restricted to the target region were used for the study. Identification of brain regions was based on anatomical landmarks and on a mouse brain atlas (Paxinos & Franklin, 2012). In retrograde tracing experiments, the distribution of HP-projecting VGLUT3 neurons and colocalization with 5-HT markers was quantified over the entire rostro-caudal axis of the raphe nuclei (B5-B9 nuclei) at 100 µm interval. Raphe subdivisions were determined based on Tph2 staining (Paxinos & Franklin, 2012). Distance from midline was analyzed using a custom MATLAB script (Mathworks, Inc).

### Statistics

All figures show mean ± SEM except for Figure 6H which shows median ± 1 standard deviation. Statistical analysis was performed using OriginLab (OriginLab Corporation), all tests used are described in the results section, p-values of < 0.05 were considered statistically significant.

## Results

### VGLUT3-positive projections from the DR and median raphe region (MRR) contact distinct brain areas

We characterized long-range projection patterns of VGLUT3-positive neurons from the DR and MnR. VGLUT3-positive projections were visualized using an anterograde Cre-dependent adeno-associated virus (AAV) expressing the membrane targeted hChR2_(E123T/T159C)_-eYFP fusion protein, injected in VGLUT3-Cre mice. Raphe subdivisions were identified using immunostaining for Tryptophane hydroxylase 2 (Tph2), the rate-limiting enzyme for 5-HT synthesis. Specificity of Cre expression in VGLUT3-Cre mice was validated using double-fluorescent *in situ* hybridization detection of VGLUT3 and Cre mRNAs along the entire rostro-caudal axis of the B5-B9 raphe nuclei. In the DR and MnR respectively, 99.48% and 99.27% of VGLUT3-positive neurons co-expressed Cre recombinase (Figure 1A, n = 3 mice, 972 neurons in DR, 411 neurons in MnR). All Cre-positive neurons expressed VGLUT3 mRNA, demonstrating a high specificity of Cre expression.

Qualitative analysis of DR VGLUT3-positive fiber distribution was performed on n = 3 mice (Figure 1B-C). Absence of virus expression in the MnR was carefully verified across the rostro-caudal extent of the raphe for each animal (Figure 1C) and the similarity of fiber expression patterns was assessed across all mice injected in the DR. In the olfactory bulb, we observed a high density of eYFP-positive fibers in the granular cell layer (Figure 1D). In the somatosensory cortex, DR VGLUT3-positive fibers were found predominantly in Layers 1 and 4 along the entire rostro-caudal axis (Figure 1E-1). A high density of VGLUT3-positive fibers from the DR was also found in the dorsal part of the claustrum and Layer 3 of the piriform cortex (Figure 1E-2, E-3). In the amygdaloid complex, we noted dense glutamatergic innervation of the central amygdala and sparse innervation of the basolateral amygdala, with very few fibers in other amygdala sub-nuclei (Figure 1F-1), as described previously (Sengupta et al., 2017). Layers 1 and 6 of the ectorhinal cortex also showed a high density of VGLUT3-positive fibers (Figure 1F-2). In the hypothalamus, a high density of fibers was observed in the parasubthalamic nucleus and the most lateral part of the lateral hypothalamic area (Figure 1F-3). As expected (Geisler et al., 2007; Qi et al., 2014), dense glutamatergic inputs from the DR were observed in the ventral tegmental area, with some fibers also present in the substantia nigra, pars compacta and the medio-dorsal part of pars reticulata (Figure 1G). We observed little to no VGLUT3-positive fibers originating from the DR in the hippocampal formation (dentate gyrus, hippocampus proper, subiculum and entorhinal cortex, data not shown).

In contrast, local injection in the MnR of an anterograde Cre-dependent AAV expressing eYFP (Figure 2A, n = 2 mice) revealed VGLUT3 neuron projections mainly to midline regions (Figure 2D-G). Accuracy of our injection sites and absence of eYFP-positive neurons in the DR were assessed along the full rostro-caudal axis of the raphe nuclei for both mice (Figure 2B). We observed a pattern of fiber expression that was mostly non-overlapping with DR VGLUT3 projections. eYFP expression was absent from the olfactory bulb and somatosensory, piriform and ectorhinal cortices (data not shown). Glutamatergic fibers from the MnR were found throughout the entire antero-posterior axis of the motor cortex with highest expression in the Layers 2 and 3; sparse expression was also observed in the anterior cingulate area (Figure 2C). Glutamatergic neurons from the MnR have been shown to project to the prefrontal cortex (Szonyi et al., 2016); accordingly, both prelimbic and infralimbic regions showed high fiber density in Layer 5 with sparse expression in other layers (Figure 2C-2). Glutamatergic fibers were also densely present in the Layer 3 of the taenia tecta (Figure 2C). One of the highest densities of MnR VGLUT3-positive fibers was observed in the septal area (Figure 2D). In the lateral septum, we noted a dense plexus of glutamatergic fibers, with some of them forming pericellular baskets closely surrounding somas and proximal dendrites of a subset of lateral septum neurons (Figure 2D-1); VGLUT3-positive neurons from the MnR also heavily targeted the medial septum (Figure 2D-2) and the diagonal band of Broca (Figure 2D-3), confirming previous observations (Jackson et al., 2009; Senft et al., 2021; Szonyi et al., 2016). In the claustrum, MnR glutamatergic projections were found mainly in the ventral part (Figure 2D-4). In the amygdaloid complex, VGLUT3-positive fibers from the MnR were most present in the lateral amygdala, sparse in the basolateral amygdala and almost completely absent in the central amygdala (Figure 2E-1). Dense fiber coverage was also observed in the medial part of the lateral hypothalamic area, surrounding the fornix (Figure 2E-2). In contrast with VGLUT3 projections from the DR that were restricted to the dorso-lateral part of the lateral hypothalamic area, MnR inputs were mainly found ventro-medially surrounding the fornix, suggesting a regional organization of VGLUT3-positive raphe inputs to the lateral hypothalamus. In contrast with DR projections, the ventral tegmental area contained very few inputs from MnR VGLUT3-positive neurons while dense expression was present in the supramammillary nucleus (Figure 2F). In the interpeduncular nucleus we also found a high density of glutamatergic fibers from the MnR (Figure 2G), but none from the DR (data not shown). We observed MnR VGLUT3-positive projections in the dorsal and lateral parts of the DR (Figure 2H). In contrast, DR VGLUT3 neurons did not send projections to the MnR (data not shown).

Finally, we analyzed the distribution of MnR VGLUT3-positive fibers in the hippocampal formation. Our AAV injection in the MnR resulted in dense innervation of the dHP, as expected based on previous studies (Jackson et al., 2009; Szonyi et al., 2016). In addition, we observed projections to the vHP, subiculum and entorhinal cortex (Figure 3). In CA1, fibers were concentrated at the junction between stratum radiatum and stratum lacunosum-moleculare, sparse in stratum oriens (Or) and absent from pyramidal layer (Pyr) (Figure 3A-1 and B-2). In CA3, stratum radiatum layer was densely packed with glutamatergic fibers (Figure 3A-2 and B-3). In the dentate gyrus, a high density of VGLUT3 MnR fibers was found in the hilus (Figure 3A-3 and B-4). In the subiculum, VGLUT3-positive fibers from the MnR were found predominantly in the molecular layer (Figure 3B-1). For CA1, CA3, dentate gyrus and subiculum, fibers distribution was similar across the dorsal and ventral regions. Finally, in the entorhinal cortex, MnR VGLUT3 fiber density was high in the lateral entorhinal cortex and sparse in the medial entorhinal cortex (Figure 3B-5). In the hippocampal formation (Figure 3C-F) and all other regions examined, eYFP and VGLUT3 staining showed a high degree of colocalization, as expected for AAV-driven eYFP expression in raphe VGLUT3-positive fibers.

Colocalization between VGLUT3 and serotonergic markers varies across different raphe subregions, for example, the most ventral part of the DR exhibits strong overlap between markers while VGLUT3 and 5-HT populations are mostly segregated in the MnR (Hioki et al., 2010; Okaty, Commons, & Dymecki, 2019; Ren et al., 2019; Sos et al., 2017). Therefore, we qualitatively examined the colocalization of eYFP in raphe-originating VGLUT3-positive fibers with 5-HT in the forebrain. eYFP-positive fibers arising from the DR frequently coexpressed 5-HT, as shown for example in somatosensory (Figure 4A) and piriform (Figure 4B) cortices, central amygdala (Figure 4C) and ventral tegmental area (Figure 4D). On the other hand, eYFP-positive fibers from the MnR rarely colocalized with 5-HT, as shown in the motor cortex (Figure 4E), dorsolateral entorhinal cortex (Figure 4F), lateral amygdala (Figure 4G) and diagonal band of Broca (Figure 4H). Overall, the amount of 5-HT co-expression in eYFP-positive fibers from the raphe nuclei generally followed the levels of co-expression between VGLUT3 and 5-HT at somatic level. DR-originating fibers showed high levels of colocalization with 5-HT while MnR-originating fibers showed little.

Collectively, these results identify brain regions with predominant input from either DR VGLUT3-positive neurons (olfactory bulb, somatosensory cortex and ventral tegmental area) or MnR VGLUT3-positive neurons (prefrontal cortex, motor cortex, septal area, supramammillary, interpeduncular and hippocampal formation). These tracing experiments also suggest a contrasted regional topography of DR versus MnR VGLUT3-positive inputs in the claustrum, amygdaloid complex and the lateral hypothalamic area. Overall, raphe VGLUT3 projection to the forebrain align with previously described pattern of raphe projections, with DR glutamatergic inputs densely targeting cortical areas (ectorhinal, pirifom, somato-sensory), dopamine rich regions (VTA and SN pars compacta), and hypothalamic areas, while MnR glutamatergic inputs heavily target midline regions (medial septum, diagonal band of Broca, lateral septum, supramammillary and interpeduncular nucleus), some cortical areas (motor, prelimbic cortex and cingulate) and the hippocampal formation (Muzerelle et al., 2014; Vertes & Linley, 2008; Xu et al., 2021). A handful of regions (amygdaloid complex, claustrum and hypothalamus) receive inputs from both the DR and MnR that appear locally segregated.

### HP-projecting VGLUT3 neurons are mainly non serotonergic and distribute over multiple raphe sub-nuclei

As shown in Figure 3, small volume injections restricted to VGLUT3-positive neurons in the MnR showed layered glutamatergic innervation along the entire dorso-ventral axis of the dentate gyrus and hippocampus proper (CA1-CA3). Yet, other raphe sub-nuclei such as B9 have also been shown to send projections to the HP in mice (Muzerelle et al., 2014). Therefore, we analyzed the distribution of VGLUT3^HP^ neurons within the raphe sub-nuclei using a retrograde viral vector approach.

The retrograde Cre-dependent viral vector AAVretro-Ef1a-DIO-eYFP was injected bilaterally in the dHP and vHP to cover the full longitudinal axis (Figure 5A). Retrograde expression of eYFP in VGLUT3 neurons was quantified along the full rostro-caudal axis of the raphe nuclei (Figure 5B). In addition, we quantified the proportion of mixed glutamatergic-serotonergic neurons within VGLUT3^HP^ population using the serotonergic marker Tph2 (Figure 5B-D). Overall, VGLUT3^HP^ neurons were distributed among several raphe sub-nuclei, with the highest densities found in B9, median (MnR) and paramedian (PMnR) raphe regions (Figure 5C). Smaller numbers of VGLUT3^HP^ neurons were also found in the interpeduncular (IP), the oral part of the pontine reticular nucleus (PnO) and the interfascicular part of the dorsal raphe (DRI). DR ventral (DRV), dorsal (DRD) and lateral (DRL) sub-regions and pontine raphe nucleus (PnR) contained little or no eYFP-positive somas (Figure 5C). Of all identified VGLUT3^HP^ neurons, a large majority (75.4 ± 2.2%, n = 6 mice) were purely glutamatergic, i.e. lacked expression of Tph2. Most HP-projecting ‘pure’ glutamatergic neurons were located in the MnR (26.9 ± 2.2%) and PMnR (20.0 ± 1.1 %). The B9 nucleus (13.9 ± 1.9%), interpeduncular (IP, 7.2 ± 2.3%) and PnO (5.6 ± 1.7%) also contained a modest population of non-serotonergic VGLUT3^HP^ neurons, while DRD (0.1 ± 0.1%), DRV (0.1 ± 0.0%), DRC (0.2 ± 0.1%), DRL (0.0 ± 0.0%) and PnR (0.9 ± 0.2%) contained very little or none (Figure 5C). Co-expression of serotonergic markers was generally low: among all VGLUT3^HP^ neurons, 24.6 ± 2.2% co-localized with Tph2. We found the highest proportion of Tph2-positive VGLUT3 neurons in the DRI (5.4 ± 1.1%), MnR (5.8 ± 0.7%) and PMnR (5.6 ± 0.9%) (Figure 5C, n = 6 mice). Other raphe subnuclei contained very few VGLUT3/Tph2 neurons (IP: 0.7 ± 0.4%; B9: 1.9 ± 0.4%, PnO: 1.7 ± 0.3%; DRD: 0.3 ± 0.1%: DRV: 2.0 ± 0.4%; DRC: 1.2 ± 0.4%; DRL and PnR: 0.0 ± 0.0%). The rostro-caudal distribution of VGLUT3^HP^ neurons showed that non-serotonergic VGLUT3^HP^ neurons were located predominantly in the rostral half of the raphe nuclei between -3.9 and -4.5 mm from bregma (Figure 5D, green, n = 6 mice). In contrast, Tph2-positive VGLUT3^HP^ neurons were evenly distributed along the rostro-caudal axis of the raphe nuclei (Figure 5D, grey).

Taken together, retrograde tracing revealed that VGLUT3^HP^ neurons distribute over several raphe subnuclei in the most rostral half of the raphe, and are predominantly found in B9, MnR and PMnR nuclei. In addition, VGLUT3^HP^ neurons are in majority purely glutamatergic, i.e. lack co-expression of the 5-HT biosynthesis enzyme Tph2.

### Distinct glutamatergic populations target the dHP and vHP, unilaterally

The dorsal and ventral poles of the HP are functionally distinct. Therefore, we sought to determine if distinct or overlapping VGLUT3^HP^ neuron populations contact the dHP and vHP. VGLUT3-Cre mice were injected with AAVretro-CAG-FLEX-tdTOM and AAVretro-Ef1a-DIO-eYFP in the dHP and vHP, bilaterally (Figure 6A and B). Proportion of VGLUT3-positive neurons projecting to the dHP, vHP or both were quantified in coronal sections of the raphe nuclei (Figure 6C). We found that among all VGLUT3^HP^ neurons, 42% projected to the dHP, 43% projected to the vHP while only 15% projected to both (Figure 6D, 1648 neurons, n = 4). In mice injected for this experiment, we sometimes noted a limited overlap in viral vector expression, revealed by co-expression of tdTOM and eYFP in VGLUT3-positive interneurons in the HP. To evaluate the contribution of this overlap to the number of VGLUT3-positive neurons projecting to both dHP and vHP, we injected mice with smaller volumes of viral vectors. In these conditions (‘small volume injections’, Figure 6E, 599 neurons, n = 2), 52% of VGLUT3^HP^ neurons projected to the dHP, 41% projected to the vHP while only 7% projected to both parts of the HP. We then analyzed the distribution of VGLUT3 neurons projecting to the dHP, vHP or both within each raphe subnucleus. Similar proportions of VGLUT3 neurons targeted the dHP versus vHP in B9 nucleus (6.3 ± 0.9% vs 4.6 ± 1.2%), the MnR (14.0 ± 1.5% vs 16.1 ± 2.6%), the PMnR (11.6 ± 0.8% vs 8.7 ± 1.0%) and in all DR sub-regions (PnR: 0.7 ± 0.2% vs 0.6 ± 0.2%; DRD: 0.0 ± 0.0% vs 0.1 ± 0.1%; DRV: 0.5 ± 0.3% vs 0.8 ± 0.4%; DRI: 2.5 ± 0.5% vs 1.9 ± 0.5%; DRC: 0.3 ± 0.2% vs 0.8 ± 0.5%, Figure 6F, n = 6 mice, Mann-Whitney test *p* > 0.05 for all regions cited above). Unbalanced dHP versus vHP innervation was observed for the IP, preferentially targeting the vHP (dHP: 1.9 ± 0.6% vs vHP: 6.8 ± 1.8%, Figure 6F, n = 6, Mann-Whitney test, *p* = 0.045) and the PnO, preferentially targeting the dHP (dHP: 7.1 ± 2.1% vs vHP: 1.8 ± 0.3%, Figure 6F, n = 6, Mann-Whitney test, *p* = 0.045).

The MRR, corresponding to the MnR, PMnR and PnO, contains almost 70% of all VGLUT3^HP^ neurons (see Figure 6F). Within the MRR, we observed that dHP-projecting somas seemed frequently located laterally while vHP-projecting somas seemed closer to midline (see example in Figure 6C). We therefore quantified the medio-lateral and rostro-caudal distributions of dHP-projecting and vHP-projecting somas in the MRR (analysis was restricted to MnR, PMnR and PnO within -4.0 mm to -4.5 mm, Figure 6H and I). First, we found that within the MRR, dHP- and vHP-projecting VGLUT3 neurons follow a distinct rostro-caudal distribution. Indeed, vHP-projecting neurons are located predominantly in the most rostral part of the MRR while dHP-projecting VGLUT3 neurons were more evenly distributed across the full rostro-caudal axis (Figure 6G). In addition, we confirmed a skewed distribution of dHP- and vHP-targeting neurons relative to midline. Distance from midline was compared for each individual soma at different raphe levels (Figure 6H). Between -4.0 and -4.2 mm from bregma, MRR VGLUT3 neurons projecting to the dHP are located significantly more laterally than vHP-projecting neurons (data is reported as mean ± SD, dHP vs vHP, n = 41-128 neurons from n = 6 mice, Mann-Whitney test; at -4.0 mm: 162.7 ± 149.9 vs 62.9 ± 77.9, *p* = 4.25E-4; at -4.1 mm: 152.3 ± 141.0 vs 70.2 ± 97.6, *p* = 9.22E-6; at -4.2 mm: 129.0 ± 144.6 vs 58.0 ± 79.2, *p* = 2.91E-5). At more caudal levels, dHP- and vHP-projecting VGLUT3 neurons were distributed similarly across the medio-lateral axis (n = 19-112 neurons from n = 6 mice; at -4.3 mm: 106.8 ± 110.8 vs 90.8 ± 98.1, *p* = 0.041; at -4.4 mm: 86.5 ± 102.1 vs 73.2 ± 65.8, *p* = 0.96; at -4.5 mm: 81.6 ± 91.1 vs 69.0 ± 60.2, *p* = 0.96). Taken together, these results show that within the MRR, VGLUT3 neurons projecting to the dHP and vHP are spatially segregated along the medio-lateral and rostro-caudal axes. We verified that the low level of overlap between dHP- and vHP-projecting VGLUT3 neurons was not due to mutually exclusive expression of the viral tracers by co-injecting both retrograde viruses in the dHP (Figure 6J-K). Among all labelled neurons, 33% expressed eYFP only, 8% expressed tdTOM only while 59% expressed both eYFP and tdTOM (Figure 6L-M, 263 neurons, n = 2 mice), confirming that both retrograde AAVs can be co-expressed in a high proportion of neurons. In addition, we controlled for non-specific retrograde fluorophore expression by injecting C57BL6/J mice in the whole hippocampus with either of the retrograde viral vectors (n = 2). We observed no eYFP-positive or tdTOM-positive somas in the raphe nuclei of injected mice (data not shown).

Finally, we examined if VGLUT3^HP^ neurons projected unilaterally or bilaterally. Mice were injected with two Cre-dependent retrograde viral vectors expressing different fluorescent proteins with one virus injected in the left HP while the other was injected in the right HP, each one covering most of the dorso-ventral axis (Figure 7A-B). Only 4% of all labelled neurons projected bilaterally to both hippocampi while 51% projected to the left and 45% to the right HP (Figure 7C-D, 426 neurons, n = 2 mice).

**Figure 7.**
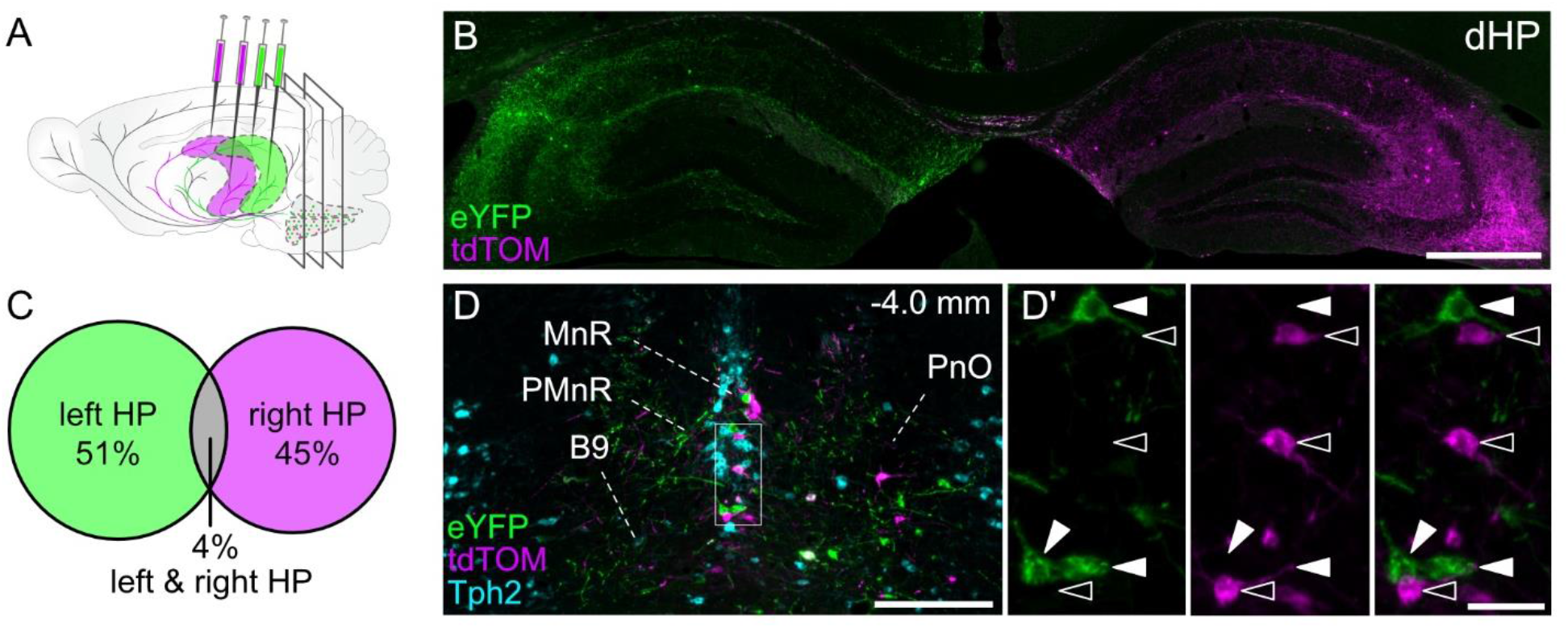
VGLUT3^HP^ neurons project unilaterally. (**A**) Schematic of two-color Cre-dependent retrograde viral vector injections in the left and right HP of VGLUT3-Cre mice. (**B**) Representative eYFP (green) and tdTOM (magenta) expression in the dHP. (**C**) Venn diagram showing that VGLUT3^HP^ neurons project mainly unilaterally with only 4% of overlap (426 neurons, n = 2 mice). (**D**) Representative eYFP (green) and tdTOM (magenta) retrograde viral expression in the MRR at -4.0 mm from bregma. Raphe subregions were identified using Tph2 immunofluorescence (cyan). Inset in (D’) shows left-HP projecting VGLUT3+ neurons (full arrows) and right-HP projecting VGLUT3+ neurons (empty arrows). Scale bars: 500 µm in B, 250 µm in D and 50 µm in D’.

Our results uncover largely independent streams of glutamatergic VGLUT3-positive inputs from the raphe to the dorsal and ventral sub-regions of the HP, in each hemisphere. In addition, we show that in the rostral part of the MRR, dHP- and vHP-targeting VGLUT3 neurons are spatially organized relative to midline, with vHP projecting cells located more medially, and dHP projecting cells more laterally. Taken together, these data suggest differential modulation of functionally distinct HP compartments by the raphe.

## Discussion

In this study, we used conditional tracing methods to target VGLUT3-positive neurons in the raphe nuclei and anatomically characterize their projection patterns to the forebrain. Systematic comparison of DR and MnR glutamatergic projections highlighted largely non-overlapping target regions, in line with the differential roles of raphe subdivisions (Ohmura et al., 2019; Teissier et al., 2015). In particular, the HP is targeted by two distinct streams of MnR VGLUT3 neurons, that innervate the dorsal and ventral poles independently. Further analysis revealed a topographic organization of VGLUT3^HP^ neurons within raphe subdivisions along the rostro-caudal and mediolateral axis. Taken together, our anatomical observations describe parallel pathways for modulation of defined target regions (or sub-regions) and functions.

Our anterograde tracing uncovered highly complementary projection patterns of VGLUT3-positive neurons originating in the DR and MnR. These findings align with cell-type independent tracing studies describing the general organization of MnR versus DR projections (Vertes & Linley, 2008) and projections of 5-HT neurons (Muzerelle et al., 2014) and VGLUT2 neurons (Szonyi et al., 2019; Xu et al., 2021). In addition, we confirmed results from several studies describing functional inputs from VGLUT3 DR neurons to the VTA (Liu et al., 2014; McDevitt et al., 2014; Qi et al., 2014; H. L. Wang et al., 2019; Zou et al., 2020) and amygdala (Sengupta et al., 2017; Sengupta & Holmes, 2019) and inputs from the MnR to the dorsal HP (He et al., 2022; Varga et al., 2009). To be noted, a very small AAV volume (25-35 nL) was injected in the MnR to completely avoid leakage into the DR, potentially leading to partial labelling of VGLUT3-positive fibers. However, in the animals used in this experiment, targeted regions were identical and only the fiber density varied. Similarly, for anterograde tracing of VGLUT3 DR fibers, two injection sites covering most of the rostro-caudal axis resulted in identical projection patterns in all three mice.

We observed a limited number of regions receiving VGLUT3-positive glutamatergic inputs from both the DR and MnR, among them the claustrum, the amygdaloid complex and hypothalamus (see Figures 1 and 2). Within these regions, DR and MnR inputs appear mostly segregated. Indeed, the DR targets preferentially the dorsal part of the claustrum while MnR VGLUT3 neurons send inputs mainly to the ventral claustrum. Similarly, dense projections from DR VGLUT3 neurons are found in the central amygdala while the MnR targets principally the lateral amygdala. In the hypothalamus, DR VGLUT3 neurons target the parasubthalamic nucleus and the lateral part of the lateral hypothalamic area while MnR-originating fibers are located more medially, surrounding the fornix. Therefore, functionally distinct subdivisions of claustrum, amygdaloid complex and hypothalamus could be differentially modulated by glutamatergic inputs from the DR and MnR.

We also observed projections between the DR and the MnR. Glutamatergic fibers from the MnR were present in the DR, but no fibers from the DR were seen in the MnR. Similarly, DR was shown to receive 5-HT inputs from the MnR, but not the opposite (Pollak Dorocic et al., 2014), suggesting mostly unidirectional modulation between these two raphe sub-nuclei. Recent studies have described inputs to identified neuronal populations in the DR and MnR (see (Pollak Dorocic et al., 2014) for 5-HT, (Xu et al., 2021) for VGLUT2 and GABA), yet no study to date has investigated inputs to DR and MnR VGLUT3 neurons. Future results describing inputs to VGLUT3 neurons would complete the overview of the anatomical properties of raphe VGLUT3 glutamatergic circuits and bring insight into their function.

Co-localization with 5-HT can be observed in both DR- and MnR-originating VGLUT3 fibers, albeit in variable proportions (Figure 4). In the DR, a large population of VGLUT3-positive neurons co-express serotonergic markers (Hioki et al., 2010; Ren et al., 2019). As expected, we observed overlap between the projection patterns of DR 5-HT neurons (Muzerelle et al., 2014) and VGLUT3 neurons (Figure 1), and frequent co-expression of 5-HT in eYFP-positive fibers from the DR (Figure 4). Interestingly, a recent study describing segregated outputs from purely 5-HT neurons compared to mixed 5-HT/VGLUT3 neurons suggests that neurochemically distinct VGLUT3 DR populations contact distinct targets (Ren et al., 2019). In the MnR, 5-HT and VGLUT3 neurons have been described as mostly non-overlapping (Sos et al., 2017). Accordingly, 5-HT co-labelling in MnR VGLUT3 fibers was overall sparse (Figure 4). Yet, we observed some similarities between the target regions of 5-HT neurons (Muzerelle et al., 2014) and VGLUT3 of the MnR (for example in the HP, lateral septum or interpeduncular nucleus), suggesting that largely non-overlapping 5-HT and VGLUT3 long-range projecting MnR neurons share common forebrain targets. Despite a growing number of studies investigating the contribution of glutamate/5-HT co-transmission, in particular in the DR (Luo, Zhou, & Liu, 2015; Sengupta et al., 2017; H. L. Wang et al., 2019; Zou et al., 2020), further investigation using intersectional strategies, targeting purely 5-HT, purely glutamatergic and mixed populations (Ren et al., 2019) are needed to uncover the fine topography of DR and MnR pathways and their functional significance.

Our retrograde tracing uncovered a distribution of VGLUT3^HP^ neurons over multiple raphe subregions. We show that in addition to MnR and PMnR, the B9 nucleus contains a large population of VGLUT3^HP^ neurons. Although less studied than other raphe subregions, B9 has been previously shown to project to the HP (Kohler & Steinbusch, 1982; Muzerelle et al., 2014) and could be involved in modulating HP function. Smaller populations of VGLUT3^HP^ neurons were also found in the IP, PnO and DRI. Therefore, with the exception of the DRI, VGLUT3-positive projections to the HP arise almost exclusively from the ventral part of the raphe in mice, similarly to what has been described for 5-HT projections (see (Muzerelle et al., 2014)). Interestingly, 5-HT neurons of the DRI and DRC, collectively forming the B6 group, have developmental origins and connectivity patterns that are closer to the MnR than the DR (Commons, 2015; Jacobs & Azmitia, 1992). The similarity in projection targets of VGLUT3-positive neurons in the DRI and MnR supports the idea that despites its ‘dorsal’ label, the DRI is more closely related to ventral sub-regions of the raphe (Commons, 2020).

The proportion of VGLUT3 neurons targeting the dHP and vHP was equivalent in most raphe subregions except for the IP, preferentially projecting to the vHP and the PnO, heavily targeting dHP (Figure 5F). In addition, our results uncover a spatial organization of VGLUT3^HP^ neurons across the mediolateral and rostro-caudal axis of the raphe. vHP-projecting VGLUT3 neurons are mainly located close to midline in the most rostral part of the raphe nuclei, whereas dHP-projecting VGLUT3 neurons are distributed evenly relative to midline and across the rostro caudal axis. This three-dimensional organization is reflected in the imbalance of dHP-versus vHP-projecting VGLUT3 neurons in the PnO (one of the most lateral raphe subregions) and the IP (one of the most rostral subregions). IP 5-HT neurons have been shown to project to the vHP in rats (Wirtshafter, Asin, & Lorens, 1986) and more recently in mice (Sherafat et al., 2020), and suggested to play a role in active stress coping and natural reward. VGLUT3-positive projections to the vHP therefore constitute a parallel IP-vHP pathway that could also play an important role in regulating vHP function. Taken together, our results suggest that glutamatergic long-range projections to the forebrain follow a fine topographical organization that awaits further investigation.

We described distinct raphe glutamatergic populations targeting the dHP and vHP in each hemisphere, suggesting independent modulation of each region and associated functions. Accordingly, dHP projecting glutamatergic neurons are likely to contribute to spatial learning and memory while vHP projecting glutamatergic neurons are likely to contribute to anxiety, social memory, and attention. In parallel, raphe-HP glutamatergic pathways are well positioned to modulate hippocampal rhythms, a role that has been suggested by several studies (Crooks et al., 2012; Domonkos et al., 2016; W. Huang et al., 2022; Jelitai, Barth, Komlosi, Freund, & Varga, 2021; D. V. Wang et al., 2015). Our anterograde tracing describes a layered organization of VGLUT3-positive fibers along the longitudinal axis of the HP, with a high density in CA1-CA3 at the junction between stratum radiatum and stratum lacunosum-moleculare. This area hosts somas of several classes of interneurons (Pelkey et al., 2017), and glutamatergic inputs from the MnR have been shown to activate some interneuron subtypes, including cholecystokinin-positive interneurons (Varga et al., 2009). Interneurons are key players in generation and/or modulation of HP rhythms such as theta (Amilhon et al., 2015), gamma (Buzsaki & Wang, 2012) and sharp wave ripples (Stark et al., 2014). Fast activation of specific classes of interneurons by VGLUT3-positive inputs from the raphe nuclei is an attractive hypothesis to explain the powerful modulation of HP rhythms by the MnR (Jackson, Dickson, & Bland, 2008; Vertes, 1981; D. V. Wang et al., 2015). Alternatively, VGLUT3-positive raphe inputs could modulate HP activity indirectly through the medial septum-diagonal band of Broca (MS-DBB) region, as suggested for 5-HT (Vertes, Hoover, & Viana Di Prisco, 2004) and VGLUT2-positive MRR inputs (Szonyi et al., 2019). Taken together, the raphe nuclei, and in particular the MnR, host multiple parallel pathways targeting the HP (5-HT and VGLUT3) or HP-related regions such as MS-DBB (5-HT, VGLUT3, VGLUT2 and GABA). The cellular targets of these distinct ascending pathways have been partially described (Chittajallu et al., 2013; Szonyi et al., 2019; Varga et al., 2009), along with local connectivity within the raphe (W. Huang et al., 2022; Jelitai et al., 2021). Yet, many elements are still missing to fully understand how these neurochemically, and anatomically diverse pathways act to modulate HP activity and function.

To conclude, we update the description of raphe VGLUT3-positive pathways to the forebrain by revealing highly segregated projections from the DR and MnR. In addition, we provide the first description of raphe VGLUT3-positive glutamatergic populations that target the dHP and vHP independently. The functional significance of this differential glutamatergic input onto distinct HP sub-regions awaits further investigation, but such parallel organization could be key to understand the modulation of HP function by raphe inputs.

## Abbreviations

5-HT: 5-hydroxytryptamine, serotonin
DBB: diagonal band of Broca
dHP: dorsal hippocampus
DR: dorsal raphe
DRD: dorsal part of the DR
DRI: interfascicular part of the dorsal raphe
DRL: lateral part of the DR
DRV: ventral part of the DR
HP: hippocampus
IP: interpeduncular
MnR: median raphe
MRR: median raphe region
MS: medial septum
PMnR: paramedian raphe
PnO: pontine reticular nucleus oral part
PnR: pontine raphe nucleus
TdTOM: tdTomato
Tph2: tryptophan hydroxylase 2
VGLUT2: vesicular glutamate transporter type 2
VGLUT3: vesicular glutamate transporter type 3
VGLUT3^HP^: hippocampus-projecting VGLUT3-positive neurons
vHP: ventral hippocampus

## Acknowledgments

We thank Dr Salah El Mestikawy for generously providing the VGLUT3-Cre mouse strain, Dr Sylvie Dumas (Oramacell, Paris, France) for the double-probe fluorescent *in situ* hybridisation experiment, Dr Elke Küster-Schöck from CHUSJ Research Center Plateform for Imaging by Microscopy for assistance with microscopy and Denise Carrier and the animal facility team for support in animal care. We also thank the Amilhon lab members, Jesse Jackson and Mahesh Karnani for valuable feedback and discussion of the manuscript. This work was supported by the Brain and Behavior Research Foundation (NARSAD Young Investigator Grant #25287) and the Natural Sciences and Engineering Research Council of Canada (NSERC Discovery Grant #RGPIN-2018-06765). JFH was supported by a Frederick Banting and Charles Best Canada Graduate Scholarship (Canadian Institutes of Health Research) and a scholarship from the Département de Neuroscience, Université de Montréal.

## References

Adhikari, A., Topiwala, M. A., & Gordon, J. A. (2010). Synchronized activity between the ventral hippocampus and the medial prefrontal cortex during anxiety. Neuron, 65(2), 257–269. doi:10.1016/j.neuron.2009.12.002

Amilhon, B., Huh, C. Y., Manseau, F., Ducharme, G., Nichol, H., Adamantidis, A., & Williams, S. (2015). Parvalbumin Interneurons of Hippocampus Tune Population Activity at Theta Frequency. Neuron, 86(5), 1277–1289. doi:10.1016/j.neuron.2015.05.027

Bannerman, D. M., Rawlins, J. N., McHugh, S. B., Deacon, R. M., Yee, B. K., Bast, T., … Feldon, J. (2004). Regional dissociations within the hippocampus--memory and anxiety. Neurosci Biobehav Rev, 28(3), 273–283. doi:10.1016/j.neubiorev.2004.03.004

Bannerman, D. M., Sprengel, R., Sanderson, D. J., McHugh, S. B., Rawlins, J. N., Monyer, H., & Seeburg, P. H. (2014). Hippocampal synaptic plasticity, spatial memory and anxiety. Nat Rev Neurosci, 15(3), 181–192. doi:10.1038/nrn3677

Buzsaki, G., & Wang, X. J. (2012). Mechanisms of gamma oscillations. Annu Rev Neurosci, 35, 203–225. doi:10.1146/annurev-neuro-062111-150444

Chittajallu, R., Craig, M. T., McFarland, A., Yuan, X., Gerfen, S., Tricoire, L., … McBain, C. J. (2013). Dual origins of functionally distinct O-LM interneurons revealed by differential 5-HT(3A)R expression. Nat Neurosci, 16(11), 1598–1607. doi:10.1038/nn.3538

Commons, K. G. (2015). Two major network domains in the dorsal raphe nucleus. J Comp Neurol, 523(10), 1488–1504. doi:10.1002/cne.23748

Commons, K. G. (2020). Serotonin system function, organization and feedback. In C. P. Müller & K. A. Cunningham (Eds.), Handbook of the behavioral neurobiology of serotonin, second edition (pp. 41).

Crooks, R., Jackson, J., & Bland, B. H. (2012). Dissociable pathways facilitate theta and non-theta states in the median raphe--septohippocampal circuit. Hippocampus, 22(7), 1567–1576. doi:10.1002/hipo.20999

de Lavilleon, G., Lacroix, M. M., Rondi-Reig, L., & Benchenane, K. (2015). Explicit memory creation during sleep demonstrates a causal role of place cells in navigation. Nat Neurosci, 18(4), 493–495. doi:10.1038/nn.3970

Domonkos, A., Nikitidou Ledri, L., Laszlovszky, T., Cserep, C., Borhegyi, Z., Papp, E., … Varga, V. (2016). Divergent in vivo activity of non-serotonergic and serotonergic VGluT3-neurones in the median raphe region. J Physiol, 594(13), 3775–3790. doi:10.1113/JP272036

Dumas, S., & Wallen-Mackenzie, A. (2019). Developmental Co-expression of Vglut2 and Nurr1 in a Mes-Di-Encephalic Continuum Preceeds Dopamine and Glutamate Neuron Specification. Front Cell Dev Biol, 7, 307. doi:10.3389/fcell.2019.00307

Fanselow, M. S., & Dong, H. W. (2010). Are the dorsal and ventral hippocampus functionally distinct structures? Neuron, 65(1), 7–19. doi:10.1016/j.neuron.2009.11.031

Felix-Ortiz, A. C., Beyeler, A., Seo, C., Leppla, C. A., Wildes, C. P., & Tye, K. M. (2013). BLA to vHPC inputs modulate anxiety-related behaviors. Neuron, 79(4), 658–664. doi:10.1016/j.neuron.2013.06.016

Fremeau, R. T., Jr., Burman, J., Qureshi, T., Tran, C. H., Proctor, J., Johnson, J., … Edwards, R. H. (2002). The identification of vesicular glutamate transporter 3 suggests novel modes of signaling by glutamate. Proc Natl Acad Sci U S A, 99(22), 14488–14493. doi:10.1073/pnas.222546799

Fu, W., Le Maitre, E., Fabre, V., Bernard, J. F., David Xu, Z. Q., & Hokfelt, T. (2010). Chemical neuroanatomy of the dorsal raphe nucleus and adjacent structures of the mouse brain. J Comp Neurol, 518(17), 3464–3494. doi:10.1002/cne.22407

Geisler, S., Derst, C., Veh, R. W., & Zahm, D. S. (2007). Glutamatergic afferents of the ventral tegmental area in the rat. J Neurosci, 27(21), 5730–5743. doi:10.1523/JNEUROSCI.0012-07.2007

Gras, C., Herzog, E., Bellenchi, G. C., Bernard, V., Ravassard, P., Pohl, M., … El Mestikawy, S. (2002). A third vesicular glutamate transporter expressed by cholinergic and serotoninergic neurons. J Neurosci, 22(13), 5442–5451. Retrieved from https://www.ncbi.nlm.nih.gov/pubmed/12097496

He, A., Zhang, C., Ke, X., Yi, Y., Yu, Q., Zhang, T., … Lu, Y. (2022). VGLUT3 neurons in median raphe control the efficacy of spatial memory retrieval via ETV4 regulation of VGLUT3 transcription. Sci China Life Sci. doi:10.1007/s11427-021-2047-8

Herzog, E., Gilchrist, J., Gras, C., Muzerelle, A., Ravassard, P., Giros, B., … El Mestikawy, S. (2004). Localization of VGLUT3, the vesicular glutamate transporter type 3, in the rat brain. Neuroscience, 123(4), 983–1002. doi:10.1016/j.neuroscience.2003.10.039

Hioki, H., Nakamura, H., Ma, Y. F., Konno, M., Hayakawa, T., Nakamura, K. C., … Kaneko, T. (2010). Vesicular glutamate transporter 3-expressing nonserotonergic projection neurons constitute a subregion in the rat midbrain raphe nuclei. J Comp Neurol, 518(5), 668–686. doi:10.1002/cne.22237

Huang, K. W., Ochandarena, N. E., Philson, A. C., Hyun, M., Birnbaum, J. E., Cicconet, M., & Sabatini, B. L. (2019). Molecular and anatomical organization of the dorsal raphe nucleus. Elife, 8. doi:10.7554/eLife.46464

Huang, W., Ikemoto, S., & Wang, D. V. (2022). Median Raphe Nonserotonergic Neurons Modulate Hippocampal Theta Oscillations. J Neurosci, 42(10), 1987–1998. doi:10.1523/JNEUROSCI.1536-21.2022

Jackson, J., Bland, B. H., & Antle, M. C. (2009). Nonserotonergic projection neurons in the midbrain raphe nuclei contain the vesicular glutamate transporter VGLUT3. Synapse, 63(1), 31–41. doi:10.1002/syn.20581

Jackson, J., Dickson, C. T., & Bland, B. H. (2008). Median raphe stimulation disrupts hippocampal theta via rapid inhibition and state-dependent phase reset of theta-related neural circuitry. J Neurophysiol, 99(6), 3009–3026. doi:10.1152/jn.00065.2008

Jacobs, B. L., & Azmitia, E. C. (1992). Structure and function of the brain serotonin system. Physiol Rev, 72(1), 165–229. doi:10.1152/physrev.1992.72.1.165

Jelitai, M., Barth, A. M., Komlosi, F., Freund, T. F., & Varga, V. (2021). Activity and Coupling to Hippocampal Oscillations of Median Raphe GABAergic Cells in Awake Mice. Front Neural Circuits, 15, 784034. doi:10.3389/fncir.2021.784034

Jimenez, J. C., Su, K., Goldberg, A. R., Luna, V. M., Biane, J. S., Ordek, G., … Kheirbek, M. A. (2018). Anxiety Cells in a Hippocampal-Hypothalamic Circuit. Neuron, 97(3), 670–683 e676. doi:10.1016/j.neuron.2018.01.016

Kjelstrup, K. G., Tuvnes, F. A., Steffenach, H. A., Murison, R., Moser, E. I., & Moser, M. B. (2002). Reduced fear expression after lesions of the ventral hippocampus. Proc Natl Acad Sci U S A, 99(16), 10825–10830. doi:10.1073/pnas.152112399

Kohler, C., & Steinbusch, H. (1982). Identification of serotonin and non-serotonin-containing neurons of the mid-brain raphe projecting to the entorhinal area and the hippocampal formation. A combined immunohistochemical and fluorescent retrograde tracing study in the rat brain. Neuroscience, 7(4), 951–975. Retrieved from http://www.ncbi.nlm.nih.gov/pubmed/7048127

Li, X., Chen, W., Pan, K., Li, H., Pang, P., Guo, Y., … Lu, Y. (2018). Serotonin receptor 2c-expressing cells in the ventral CA1 control attention via innervation of the Edinger-Westphal nucleus. Nat Neurosci, 21(9), 1239–1250. doi:10.1038/s41593-018-0207-0

Liu, Z., Zhou, J., Li, Y., Hu, F., Lu, Y., Ma, M., … Luo, M. (2014). Dorsal raphe neurons signal reward through 5-HT and glutamate. Neuron, 81(6), 1360–1374. doi:10.1016/j.neuron.2014.02.010

Luo, M., Zhou, J., & Liu, Z. (2015). Reward processing by the dorsal raphe nucleus: 5-HT and beyond. Learn Mem, 22(9), 452–460. doi:10.1101/lm.037317.114

McDevitt, R. A., Tiran-Cappello, A., Shen, H., Balderas, I., Britt, J. P., Marino, R. A., … Bonci, A. (2014). Serotonergic versus nonserotonergic dorsal raphe projection neurons: differential participation in reward circuitry. Cell Rep, 8(6), 1857–1869. doi:10.1016/j.celrep.2014.08.037

Moser, M. B., & Moser, E. I. (1998). Functional differentiation in the hippocampus. Hippocampus, 8(6), 608–619. doi:10.1002/(SICI)1098-1063(1998)8:6<608::AID-HIPO3>3.0.CO;2-7

Moser, M. B., Moser, E. I., Forrest, E., Andersen, P., & Morris, R. G. (1995). Spatial learning with a minislab in the dorsal hippocampus. Proc Natl Acad Sci U S A, 92(21), 9697–9701. doi:10.1073/pnas.92.21.9697

Muzerelle, A., Scotto-Lomassese, S., Bernard, J. F., Soiza-Reilly, M., & Gaspar, P. (2014). Conditional anterograde tracing reveals distinct targeting of individual serotonin cell groups (B5-B9) to the forebrain and brainstem. Brain Struct Funct. doi:10.1007/s00429-014-0924-4

Nectow, A. R., Schneeberger, M., Zhang, H., Field, B. C., Renier, N., Azevedo, E., … Friedman, J. M. (2017). Identification of a Brainstem Circuit Controlling Feeding. Cell, 170(3), 429–442 e411. doi:10.1016/j.cell.2017.06.045

Ohmura, Y., Tsutsui-Kimura, I., Sasamori, H., Nebuka, M., Nishitani, N., Tanaka, K. F., … Yoshioka, M. (2019). Different roles of distinct serotonergic pathways in anxiety-like behavior, antidepressant-like, and anti-impulsive effects. Neuropharmacology, 107703. doi:10.1016/j.neuropharm.2019.107703

Okaty, B. W., Commons, K. G., & Dymecki, S. M. (2019). Embracing diversity in the 5-HT neuronal system. Nat Rev Neurosci, 20(7), 397–424. doi:10.1038/s41583-019-0151-3

Okaty, B. W., Sturrock, N., Escobedo Lozoya, Y., Chang, Y., Senft, R. A., Lyon, K. A., … Dymecki, S. M. (2020). A single-cell transcriptomic and anatomic atlas of mouse dorsal raphe Pet1 neurons. Elife, 9. doi:10.7554/eLife.55523

Okuyama, T., Kitamura, T., Roy, D. S., Itohara, S., & Tonegawa, S. (2016). Ventral CA1 neurons store social memory. Science, 353(6307), 1536–1541. doi:10.1126/science.aaf7003

Paxinos, G., & Franklin, K. B. J. (2012). The Mouse Brain in Stereotaxic Coordinates, Fourth Edition: Academic Press.

Pelkey, K. A., Chittajallu, R., Craig, M. T., Tricoire, L., Wester, J. C., & McBain, C. J. (2017). Hippocampal GABAergic Inhibitory Interneurons. Physiol Rev, 97(4), 1619–1747. doi:10.1152/physrev.00007.2017

Pollak Dorocic, I., Furth, D., Xuan, Y., Johansson, Y., Pozzi, L., Silberberg, G., … Meletis, K. (2014). A whole-brain atlas of inputs to serotonergic neurons of the dorsal and median raphe nuclei. Neuron, 83(3), 663–678. doi:10.1016/j.neuron.2014.07.002

Pothuizen, H. H., Zhang, W. N., Jongen-Relo, A. L., Feldon, J., & Yee, B. K. (2004). Dissociation of function between the dorsal and the ventral hippocampus in spatial learning abilities of the rat: a within-subject, within-task comparison of reference and working spatial memory. The European journal of neuroscience, 19(3), 705–712. doi:10.1111/j.0953-816x.2004.03170.x

Qi, J., Zhang, S., Wang, H. L., Wang, H., de Jesus Aceves Buendia, J., Hoffman, A. F., … Morales, M. (2014). A glutamatergic reward input from the dorsal raphe to ventral tegmental area dopamine neurons. Nat Commun, 5, 5390. doi:10.1038/ncomms6390

Rao, R. P., von Heimendahl, M., Bahr, V., & Brecht, M. (2019). Neuronal Responses to Conspecifics in the Ventral CA1. Cell Rep, 27(12), 3460–3472 e3463. doi:10.1016/j.celrep.2019.05.081

Ren, J., Friedmann, D., Xiong, J., Liu, C. D., Ferguson, B. R., Weerakkody, T., … Luo, L. (2018). Anatomically Defined and Functionally Distinct Dorsal Raphe Serotonin Sub-systems. Cell. doi:10.1016/j.cell.2018.07.043

Ren, J., Isakova, A., Friedmann, D., Zeng, J., Grutzner, S. M., Pun, A., … Luo, L. (2019). Single-cell transcriptomes and whole-brain projections of serotonin neurons in the mouse dorsal and median raphe nuclei. Elife, 8. doi:10.7554/eLife.49424

Schafer, M. K., Varoqui, H., Defamie, N., Weihe, E., & Erickson, J. D. (2002). Molecular cloning and functional identification of mouse vesicular glutamate transporter 3 and its expression in subsets of novel excitatory neurons. J Biol Chem, 277(52), 50734–50748. doi:10.1074/jbc.M206738200

Senft, R. A., Freret, M. E., Sturrock, N., & Dymecki, S. M. (2021). Neurochemically and Hodologically Distinct Ascending VGLUT3 versus Serotonin Subsystems Comprise the r2-Pet1 Median Raphe. J Neurosci, 41(12), 2581–2600. doi:10.1523/JNEUROSCI.1667-20.2021

Sengupta, A., Bocchio, M., Bannerman, D. M., Sharp, T., & Capogna, M. (2017). Control of Amygdala Circuits by 5-HT Neurons via 5-HT and Glutamate Cotransmission. J Neurosci, 37(7), 1785–1796. doi:10.1523/JNEUROSCI.2238-16.2016

Sengupta, A., & Holmes, A. (2019). A Discrete Dorsal Raphe to Basal Amygdala 5-HT Circuit Calibrates Aversive Memory. Neuron, 103(3), 489–505 e487. doi:10.1016/j.neuron.2019.05.029

Sherafat, Y., Bautista, M., Fowler, J. P., Chen, E., Ahmed, A., & Fowler, C. D. (2020). The Interpeduncular-Ventral Hippocampus Pathway Mediates Active Stress Coping and Natural Reward. eNeuro, 7(6). doi:10.1523/ENEURO.0191-20.2020

Sos, K. E., Mayer, M. I., Cserep, C., Takacs, F. S., Szonyi, A., Freund, T. F., & Nyiri, G. (2017). Cellular architecture and transmitter phenotypes of neurons of the mouse median raphe region. Brain Struct Funct, 222(1), 287–299. doi:10.1007/s00429-016-1217-x

Stark, E., Roux, L., Eichler, R., Senzai, Y., Royer, S., & Buzsaki, G. (2014). Pyramidal cell-interneuron interactions underlie hippocampal ripple oscillations. Neuron, 83(2), 467–480. doi:10.1016/j.neuron.2014.06.023

Strange, B. A., Witter, M. P., Lein, E. S., & Moser, E. I. (2014). Functional organization of the hippocampal longitudinal axis. Nat Rev Neurosci, 15(10), 655–669. doi:10.1038/nrn3785

Szonyi, A., Mayer, M. I., Cserep, C., Takacs, V. T., Watanabe, M., Freund, T. F., & Nyiri, G. (2016). The ascending median raphe projections are mainly glutamatergic in the mouse forebrain. Brain Struct Funct, 221(2), 735–751. doi:10.1007/s00429-014-0935-1

Szonyi, A., Zicho, K., Barth, A. M., Gonczi, R. T., Schlingloff, D., Torok, B., … Nyiri, G. (2019). Median raphe controls acquisition of negative experience in the mouse. Science, 366(6469). doi:10.1126/science.aay8746

Teissier, A., Chemiakine, A., Inbar, B., Bagchi, S., Ray, R. S., Palmiter, R. D., … Ansorge, M. S. (2015). Activity of Raphe Serotonergic Neurons Controls Emotional Behaviors. Cell Rep, 13(9), 1965–1976. doi:10.1016/j.celrep.2015.10.061

Varga, V., Losonczy, A., Zemelman, B. V., Borhegyi, Z., Nyiri, G., Domonkos, A., … Freund, T. F. (2009). Fast synaptic subcortical control of hippocampal circuits. Science, 326(5951), 449–453. doi:10.1126/science.1178307

Vertes, R. P. (1981). An analysis of ascending brain stem systems involved in hippocampal synchronization and desynchronization. J Neurophysiol, 46(5), 1140–1159. doi:10.1152/jn.1981.46.5.1140

Vertes, R. P., Fortin, W. J., & Crane, A. M. (1999). Projections of the median raphe nucleus in the rat. J Comp Neurol, 407(4), 555–582. Retrieved from https://www.ncbi.nlm.nih.gov/pubmed/10235645

Vertes, R. P., Hoover, W. B., & Viana Di Prisco, G. (2004). Theta rhythm of the hippocampus: subcortical control and functional significance. Behav Cogn Neurosci Rev, 3(3), 173–200. doi:10.1177/1534582304273594

Vertes, R. P., & Linley, S. B. (2008). Efferent and afferent connections of the dorsal and median raphe nuclei in the rat. In J. M. Monti, S. R. Pandi-Perumal, B. L. Jacobs, & D. J. Nutt (Eds.), Serotonin and Sleep: Molecular, Functional and Clinical Aspects (pp. 69).

Wang, D. V., Yau, H. J., Broker, C. J., Tsou, J. H., Bonci, A., & Ikemoto, S. (2015). Mesopontine median raphe regulates hippocampal ripple oscillation and memory consolidation. Nat Neurosci, 18(5), 728–735. doi:10.1038/nn.3998

Wang, H. L., Zhang, S., Qi, J., Wang, H., Cachope, R., Mejias-Aponte, C. A., … Morales, M. (2019). Dorsal Raphe Dual Serotonin-Glutamate Neurons Drive Reward by Establishing Excitatory Synapses on VTA Mesoaccumbens Dopamine Neurons. Cell Rep, 26(5), 1128–1142 e1127. doi:10.1016/j.celrep.2019.01.014

Wirtshafter, D., Asin, K. E., & Lorens, S. A. (1986). Serotonin-immunoreactive projections to the hippocampus from the interpeduncular nucleus in the rat. Neurosci Lett, 64(3), 259–262. doi:10.1016/0304-3940(86)90338-1

Xu, Z., Feng, Z., Zhao, M., Sun, Q., Deng, L., Jia, X., … Li, A. (2021). Whole-brain connectivity atlas of glutamatergic and GABAergic neurons in the mouse dorsal and median raphe nuclei. Elife, 10. doi:10.7554/eLife.65502

Yoshida, K., Drew, M. R., Kono, A., Mimura, M., Takata, N., & Tanaka, K. F. (2021). Chronic social defeat stress impairs goal-directed behavior through dysregulation of ventral hippocampal activity in male mice. Neuropsychopharmacology, 46(9), 1606–1616. doi:10.1038/s41386-021-00990-y

Zou, W. J., Song, Y. L., Wu, M. Y., Chen, X. T., You, Q. L., Yang, Q., … Gao, T. M. (2020). A discrete serotonergic circuit regulates vulnerability to social stress. Nat Commun, 11(1), 4218. doi:10.1038/s41467-020-18010-w

